# Mass spectrometric identification of a PSI–LHCI–cytochrome *b*_6_*f* complex in both *Arabidopsis stn7* and *pph1*

**DOI:** 10.64898/2026.10.16.759642

**Authors:** Lutz A. Eichacker, Janine Arnold, Geoffrey Fucile, Ann Kristin Vatland, Karol Stensen, Gerhard Weber

## Abstract

Native PAGE separation of digitonin-solubilized thylakoid stroma lamellae resolves PSI and cytochrome *b*_6_*f* into separate bands. Mass spectrometric analysis of the 1D native gel bands showed that the bands are baskets of co-migrating complexes along the molecular size axis. Hierarchical clustering confirmed the separation of dimeric cytochrome *b*_6_*f* from PSI in 1D LN-PAGE. Interval-zone free-flow electrophoresis separated the same extracts by charge in a flowing buffer film, and the continuous fractionation indicated co-migration of complexes without the in-gel structural hindrances. A second-dimension LN-PAGE separated the complexes comigrating in iZE into complexes with defined molecular size, allowing in-gel mass spectrometry identification in the gel bands. Both genotypes showed a 720 kDa fluorescent band in which cytochrome *f* (PetA) and cytochrome *b*_6_ (PetB) join the PSI–LHCI node (*stn7*, *d* = 0.50; *pph1*, *d* = 0.34). Lhcb1 and Lhcb2 are absent from the 720 kDa node. The 753 kDa band is specific to *pph1* (*d* = 0.38). Phosphorylated LHCII1,2 joins PSI–LHCI there, and PetB is detected in every slice of that band without becoming a leaf of the tree.

**Highlights:**

1. A PSI complex (PSI–LHCI–cytochrome b6f) is identified by MS in both stn7 and pph1.
2. Phosphorylated LHCII1,2 binds to this PSI complex in pph1, at 753 kDa.
3. 2D iZE/LN-PAGE resolves the complex while 1D PAGE and iZE alone are insufficient.

## 1. Introduction

When the spectral composition of light changes, higher plants alter the distribution of absorbed energy between photosystem II (PSII) and photosystem I (PSI). The adjustment is known as a state transition. Reduction of the plastoquinone pool, which occurs when PSII is preferentially excited, is accompanied by phosphorylation of a fraction of the light-harvesting complex II (LHCII) and by movement of that fraction from the stacked grana membranes into the unstacked stroma lamellae, where PSI is enriched (Bennett, 1980; Allen, 1992, 2003; Rochaix, 2014). Bennett (1980) showed that the phosphorylated polypeptides are those of LHCII. In *Arabidopsis thaliana*, the fraction of the PSII antenna that takes part is about 15–20%, whereas in *Chlamydomonas reinhardtii* about 80% of LHCII participates (Allen, 1992; Delosme et al., 1996). Despite the modest size of the mobile fraction in land plants, plants lacking the responsible kinase grow poorly when light quality fluctuates, and the same kinase is required for the longer-term adjustment of photosystem stoichiometry (Bellafiore et al., 2005; Pesaresi et al., 2009; Tikkanen et al., 2010).

The kinase and the phosphatase that act on this antenna have been identified genetically, and they are distinct from the enzymes that phosphorylate the PSII core. Plastoquinol bound at the quinol-oxidation (Qo) site of the cytochrome *b*_6_*f* complex activates the LHCII kinase (Vener et al., 1997; Zito et al., 1999). In *C. reinhardtii*, the kinase is state transition 7 (Stt7); an *stt7* mutant does not phosphorylate LHCII or perform state transitions (Depège et al., 2003). The orthologous kinase of *A. thaliana* is STN7. An *stn7* null mutant lacks state transitions and LHCII phosphorylation (Bellafiore et al., 2005). The phosphatase that dephosphorylates LHCII was identified independently as the thylakoid-associated phosphatase of 38 kDa (TAP38) and as protein phosphatase 1 (PPH1). In either mutant, LHCII remains phosphorylated, and the plants remain in State 2 under conditions where the wild type would return to State 1 (Pribil et al., 2010; Shapiguzov et al., 2010). A second kinase, STN8, phosphorylates the PSII core subunits D1, D2, and CP43; an *stn8* mutant still performs state transitions (Bonardi et al., 2005). A different enzyme, the PSII core phosphatase (PBCP), dephosphorylates those core subunits. STN8 phosphorylates D1, D2 and CP43 when the leaf is illuminated, and Samol et al. (2012) showed that a *pbcp* mutant fails to remove that phosphate after the plants are returned to darkness, so the phosphorylated core subunits persist. Phosphorylation of LHCII is not the principal defect in *pbcp*. The phosphatase absent in *pph1* acts on LHCII, not on the PSII core.

Phosphoproteomic comparisons of *stn7* with the wild type assigned phosphorylation of Lhcb1 and Lhcb2, of Lhcb4 (CP29) and of Lhcb5 (CP26) to STN7, and a related analysis identified the plastid transcriptionally active chromosome protein pTAC16 as a substrate of the STN7/STN8 pair (Fristedt and Vener, 2011; Ingelsson and Vener, 2012). PPH1/TAP38 has been shown to reverse the LHCII phosphorylation lost in *stn7* (Pribil et al., 2010; Shapiguzov et al., 2010). The two mutants used in the present work therefore lack opposite activities of one regulatory sequence: *stn7* does not phosphorylate the mobile antenna, and *pph1* does not remove that phosphate.

The products of the *Lhcb* genes do not behave as a single pool during a state transition. Lhcb1 and Lhcb2 assemble the trimeric LHCII that can dissociate from PSII, and Lhcb3, although able to enter such trimers, differs from Lhcb1 and Lhcb2 in trimer formation and is present in smaller amounts (Standfuss and Kühlbrandt, 2004). Lhcb4 (CP29), Lhcb5 (CP26) and Lhcb6 (CP24) are monomeric. In the PSII supercomplexes resolved by electron microscopy, they occupy fixed positions between the reaction center and the LHCII trimers, and the peripheral trimers, rather than these monomeric subunits, are the ones lost when the supercomplex is destabilized (Dekker and Boekema, 2005; Caffarri et al., 2009). The same structural work shows a substantial amount of LHCII that is not incorporated into the C_2_S_2_M_2_ supercomplex. Describing the state transition requires keeping three populations distinct: the mobile Lhcb1/Lhcb2 trimer, the monomeric minor antenna, and LHCII not stably bound to either reaction center.

Cryo-electron microscopy has determined the site where one mobile trimer can bind PSI. The PSI–LHCI structure of pea contains the four LHCI proteins Lhca1, Lhca2, Lhca3, and Lhca4 arranged as a crescent along one side of the PSI core (Qin et al., 2015). The PSI– LHCI–LHCII supercomplex prepared from maize in State 2 also contains a single LHCII trimer bound on the opposite side of the core, in contact with PsaO, PsaH, PsaL, and PsaA (Pan et al., 2018). The Lhcb polypeptides that reach PSI in that structure are therefore attached to the reaction center and are not arranged as a further layer outside the Lhca crescent. From *C. reinhardtii*, Iwai et al. (2010) isolated a State 2 supercomplex containing PSI, LHCI, LHCII and cytochrome *b*_6_*f*, and Steinbeck et al. (2018) determined the structure of a PSI–LHCI–cytochrome *b*_6_*f* supercomplex formed under anaerobic conditions. No atomic model of cytochrome *b*_6_*f* bound to plant PSI, at a resolution comparable to the LHCII site in Pan et al. (2018), has been reported. Yadav et al. (2017) separated *pph1* thylakoids by free-flow electrophoresis and, using single-particle electron microscopy of the resulting fractions, observed particles in which PSI–LHCI contacted cytochrome *b*_6_*f*, LHCII, or the chloroplast NAD(P)H dehydrogenase-like (NDH) complex. Each study recovered a defined assembly from a defined physiological state. None establishes which fluorescent bands from native electrophoresis of *stn7* and *pph1* from plants grown under a standard light regime correspond to these assemblies.

Whether two of these proteins remain together after the membrane has been dissolved is a question of separation, and separation must be distinguished from arrangement in the intact thylakoid. In stacked grana of intact Arabidopsis leaves, the exoplasmic fracture face contains a mixture of PSII supercomplexes of different antenna sizes, including the dimeric core alone (C_2_), the core with two strongly bound LHCII trimers (C_2_S_2_), and the core with two strongly and two moderately bound trimers (C_2_S_2_M_2_), together with cytochrome *b*_6_*f*. Volume exclusion accounts for the overall clustering of these particles, whereas their local orientation indicates short-range attraction between neighbors (Mohamed and Oung, 2026). Detergent removes the lipids in which this arrangement is embedded. What remains, and what can then be separated into subclasses, are the associations that survive in the detergent micelle. At low concentration, digitonin preferentially releases the unstacked stroma lamellae and leaves PSII enriched in the granal residue (Andersson and Anderson, 1980). Co-migration after this step is evidence that the proteins stayed together once the bilayer was gone. It does not reconstruct every contact that crowding can produce in the intact membrane (Mohamed and Oung, 2026).

Native polyacrylamide electrophoresis is one way to sort the particles released by that solubilization. An electric field drives them through a continuous network whose pore size is set by acrylamide concentration and cross-linker proportion. A particle enters the gel only when its smallest cross-section is smaller than the pores at the top of the gel, and a gradient gel then retards it further as the pores narrow. The slope of the logarithm of mobility against gel concentration, the Ferguson plot, reports retardation (Ferguson, 1964). Gel-filtration chromatography uses a gel for a different purpose. There, particles diffuse into the solvent inside porous beads; no electric field is applied, and larger particles, excluded from the bead interior, travel in the void volume and leave the column first. In electrophoresis, the same particle must pass through the network, so the relevant dimension is the one that fits the pore, not the volume that excludes it from a bead.

For a spherical protein, the hydrodynamic diameter, in nanometres, is estimated from the molecular mass in daltons by 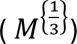. A sphere of the mass of one C_2_S_2_M_2_ supercomplex, about 1.4 MDa, would have a diameter of about 16 nm. The particle is not a sphere. Electron microscopy of membrane-bound PSII crystals from *A. thaliana* gave an in-plane unit cell of 25.6 nm by 21.4 nm for this assembly, and the complex spans only about 6 nm across the membrane (Yakushevska et al., 2001). The flat particle can pass the open mesh at the entrance of a 3–12% polyacrylamide gel. Holmes and Stellwagen (1991) estimated effective pore radii, not one diameter for a given acrylamide percentage: about 140 nm in a 4.6% gel at 1.5% cross-linker, and about 20 nm only in a 10.5% gel at 5–10% cross-linker. A gradient has no single pore size. It separates C_2_S_2_M, C_2_S_2_, and the dimeric core by increasing retardation as the mesh tightens (Caffarri et al., 2009). Apparent mass, calibrated against soluble markers, reflects this sieving, along with the detergent and lipid still bound, not the sum of the polypeptide masses (Schägger and von Jagow, 1991; Wittig et al., 2006). Blue-native PAGE and lithium dodecyl sulfate native PAGE are the two forms of this separation used for thylakoid complexes. In blue-native PAGE, the cathode buffer contains Coomassie Brilliant Blue G-250 (Schägger and von Jagow, 1991; Wittig et al., 2006). In lithium dodecyl sulfate native PAGE, the cathode buffer contains lithium dodecyl sulfate at a concentration that keeps chlorophyll bound, so PSII and PSI can be recorded in the gel by absorbance and fluorescence (Järvi et al., 2011; Arnold et al., 2014). Complexes that do not enter the polyacrylamide remain at the gel surface (Behrens et al., 2013; Eichacker et al., 2015).

Several two-dimensional arrangements separate charge and size. Charge-shift electrophoresis changes the charge of the detergent bound to a membrane protein and records the resulting mobility change, distinguishing membrane proteins from soluble ones (Helenius and Simons, 1977). Anion-exchange chromatography followed by gel filtration sorts first by charge and then by hydrodynamic volume. Blue-native PAGE followed by denaturing electrophoresis in SDS sorts native complexes and then displays their subunits. In each arrangement, the first separation reduces the mixture complexity presented to the second.

Free-flow electrophoresis separates by electrophoretic mobility, in which a buffer film carries the sample across the electric field, and the resolved zones are collected as liquid fractions (Hannig and Heidrich, 1974). Particles do not need to enter a polyacrylamide network, so material that concentrates strongly at a gel surface is only moderately focused in solution. The interval-zone form completes the separation in a few minutes, and detergent in the film keeps membrane complexes soluble (Hartmann et al., 2007; Eichacker et al., 2015; Eichacker and Weber, 2026). The fractions can be measured by spectroscopy or by mass spectrometry, or they can be applied to a native gel and sieved as described above. Free-flow electrophoresis itself is a long-established method, including the fractionation of plant organelles and membranes before proteomic analysis (Hannig and Heidrich, 1974; Barkla, 2018). Less common is interval-zone separation of native photosynthetic complexes by charge in a detergent-containing film, followed by a second native gel and mass spectrometry (Eichacker et al., 2015; Yadav et al., 2017). This sequence distinguishes a particle from complexes that share only a gel band.

Mass spectrometry of the second-dimension slices identifies a PSI–LHCI–cytochrome *b*_6_*f* complex in both *Arabidopsis stn7* and *pph1*. The experiments describe the conditions that enabled this identification. Digitonin-solubilized thylakoids of plants grown under standard short-day conditions were separated first by interval-zone free-flow electrophoresis and then by lithium dodecyl sulfate native PAGE. Because *stn7* does not phosphorylate the mobile LHCII and *pph1-3* does not remove that phosphate, the two genotypes retain the two phosphorylation states of the antenna without anaerobic incubation. On the first native gel, the genotypes differ by bands at 720 and 753 kDa, and cytochrome *b*_6_*f* is not resolved inside the high-mass PSI region. The free-flow fractions that contained these bands were applied to a second native gel. Hierarchical clustering places the 720 kDa slices of both genotypes with PSI, LHCI, and cytochrome *b*_6_*f*, with cytochrome *f* (PetA) and cytochrome *b*_6_ (PetB) joining the PSI subunits. Lhcb1 and Lhcb2 are absent from that node. Only *pph1-3* produces the 753 kDa band. Those bands are the secondary result. They contain PSI, LHCI, and the phosphorylated mobile Lhcb1 and Lhcb2 polypeptides. PetB is present in every slice of that window and is not a leaf of the tree. For an asymmetric particle like the one measured for the C_2_S_2_M_2_ supercomplex, the sieving argument predicts a difference of about 30 kDa in apparent mass if the cross-section limiting entry into the gel remains close to that of PSI and the additional LHCII trimer lengthens only one axis.

## 2. Results

### 2.1 1D LN-PAGE resolved extra bands 1, 4 and 8 in *stn7*, and extra band 7 in *pph1*

Thylakoid membranes were isolated from four-week-old *Arabidopsis thaliana* Col-0, *stn7*, and *pph1-3* plants grown in short days and adapted for 3 h at 100 µmol m⁻² s⁻¹ before harvest. The membranes were solubilized in two steps and separated by native lithium dodecyl sulfate polyacrylamide gel electrophoresis (Fig. 1A). Digitonin at 16 mM released a stroma-lamella-enriched supernatant (bands 1–11). The residual pellet, extracted with 7.2 mM digitonin plus 7.2 mM *n*-dodecyl-β-D-maltoside, gave the grana-enriched bands 12– 23.

**Figure 1.**
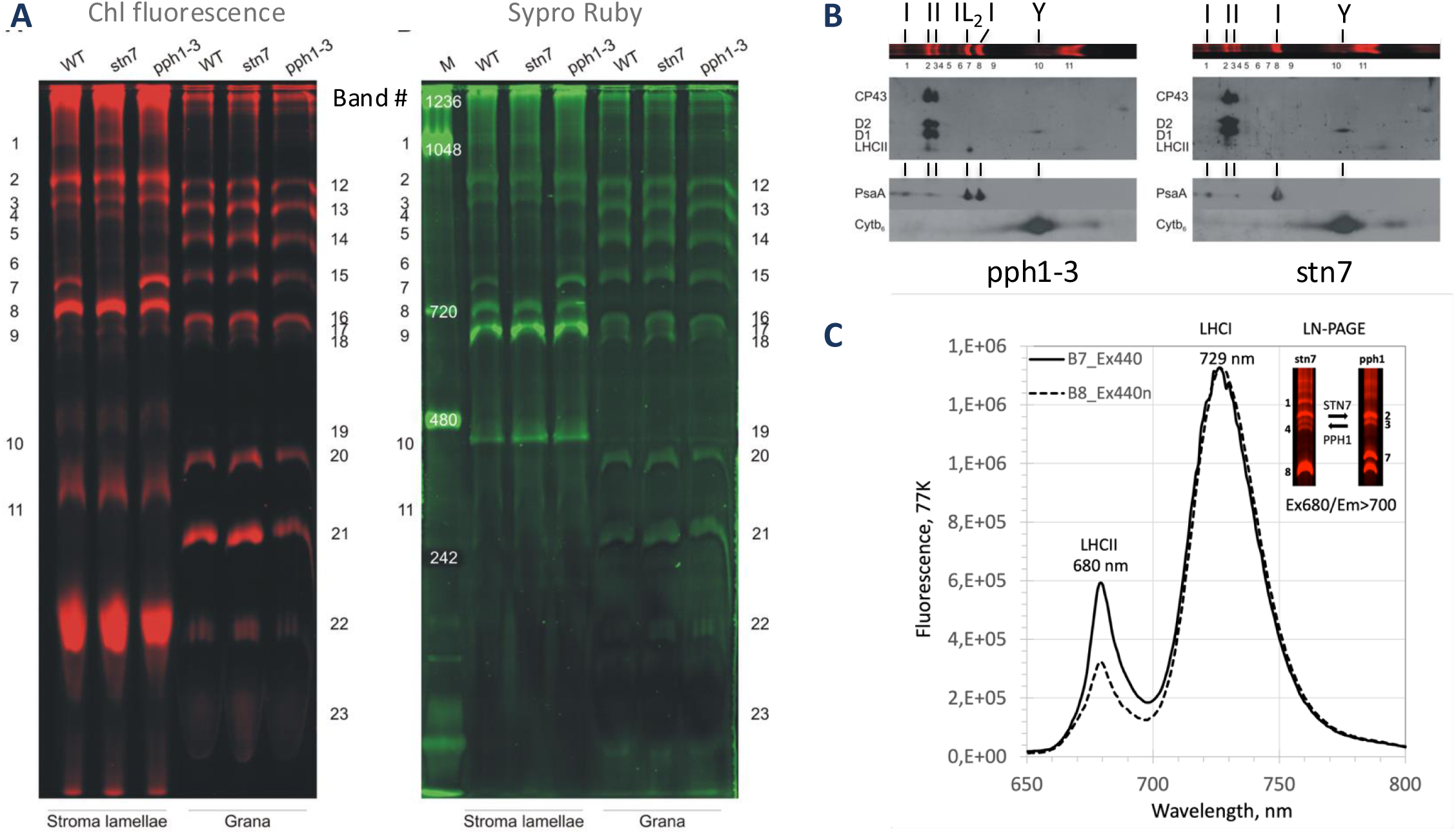
Protein band analysis of WT, stn7 and pph1 mutants after 1D- and 2D-LN-/SDS-PAGE separation. A two-step solubilization of thylakoid membranes was conducted from Arabidopsis WT, stn7, and pph1-3 mutants from “State 2”-adapted thylakoids corresponding to 8 μg Chl. Membranes were first solubilized in 16 mM digitonin (Stroma lamellae). Non-solubilized material was separated by centrifugation and was solubilized in 7.2 mM digitonin and 7.2 mM n-dodecyl β-D-maltoside (Grana). Chl-binding bands were detected by fluorescence emission scanning at 633 nm with a 670BP30 emission filter after native LDS-PAGE (A, Chl fluorescence). Total protein was stained with Sypro® Ruby and detected using an excitation wavelength of 532 nm combined with a 580BP30 filter (A, Sypro Ruby). A second-dimension SDS-PAGE of the stroma extracts (native gel lanes of stn7 and pph1-3) was wet-transferred to a nitrocellulose membrane, and the gel blots were used to identify antibodies. Antibodies against phosphothreonine identified phosphorylated PSII subunits CP43, D2, and D1 in PSII bands 2 and 3 (II) and LHCII in bands 2 and 7 (IL2). Antibodies against protein subunit PsaA identified complexes containing PSI (I) in bands 1, 2, 4, 7, 8, and antibodies against Cyt b6 identified the Cyt b6f complex (Y) in band 10. The 77 K spectra (max. 680 nm/LHCII, 731 nm/LHCI) were recorded from excised native gel slices of bands 7 (pph1) and 8 (stn7) (native gel insert Ex680/Em>700 nm), upon excitation at 440 nm (C).

We used Chl fluorescence of the gel to localize the Chl-binding photosynthetic protein complexes of the stroma and grana thylakoid membrane and applied Sypro Ruby staining of the same gel to test for the relative quantity of protein (Fig. 1A). The strongest Chl fluorescence was recorded in band 22, a band also present in the grana lane (Fig. 1A). Its protein stain was very low. The high fluorescence indicated that excitation energy was not quenched in this band, and the low protein quantity indicated that the minor LHC proteins remained largely bound to protein complexes in the higher-molecular-weight bands. The second-highest Chl fluorescence was recorded from two bands which we found to contain PSI (Fig. 1A, B). The high Chl fluorescence from these two bands was unexpected because, at RT, excitation energy at PSI is typically effectively quenched. The 77 K emission spectra then showed that both bands 7 and 8 contain LHCI (731 nm) and an LHCII emission (680 nm), which was increased in band 7 by about 1.9-fold relative to band 8 (Fig. 1B and C). Gel blot analysis confirmed that both bands contained PSI, but mobile LHCII1,2 proteins were detected only in band 7 (Fig. 1B and S2A). This difference in band structure was directly associated with the pph1 mutant (Fig. 1A). Bands 7 and 8 were present in *pph1-3* and in the wild type, while only band 8 was present in *stn7*. The next-highest fluorescence signal was from band 2, which gel-blot and MS analysis identified as a PSII-LHCII complex (Fig. 1B and 2A and B). Except for band 22, protein and fluorescence intensity appeared correlated in both membrane fractions throughout the protein complex bands, and only two protein complexes were detected by Sypro Ruby but showed no fluorescence: ATP synthase at band 9 and the cytochrome *b*_6_*f* dimer at band 10 (Fig. 1A). We further differentiated the two mutant protein complexes by analysis of bands 1 and 4, which are present in *stn7* and absent from *pph1-3*. In contrast, bands 2, 3, 6, 9, 10, and 11 were present in both mutants. To determine the molecular size of the protein complexes identified in WT and mutant plant thylakoids, we estimated band masses by comparing migration to the NativeMark standard on this gel. This analysis placed band 8 at 720 kDa and band 7 at 753 kDa.

**Figure 2A.**
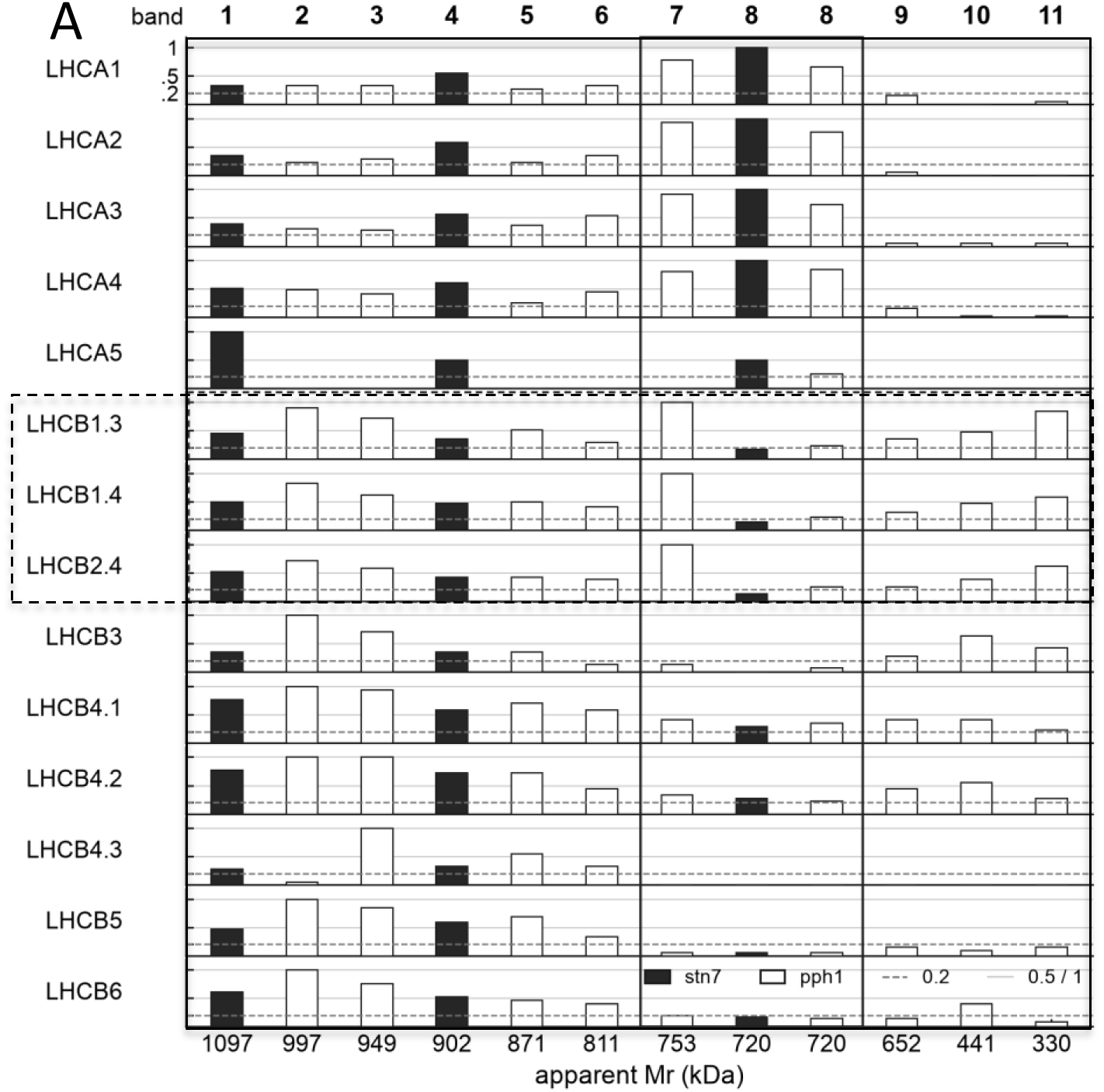
Lhc proteins identified by MS in gel-bands after 1D LN-PAGE separation of thylakoid stroma lamellae. Fluorescent protein bands were extracted after LN-PAGE, and gel slices were processed for mass spectrometric analysis. Peptides were identified by database searching (Mascot against TAIR10) and unique-peptide counts per protein were recorded. Peptide counts were normalized to the maximum unique peptide count per protein and gel slice. The corresponding peptide count distributions per LHC protein were plotted over all bands as a sparkline, as shown in the bars (max-norm). Numbers on top are gel bands. Numbers on the bottom are apparent Mr (kDa) from Figure 1A. Filled bars = stn7, open bars = pph1. Solid grey lines at 0.5 and 1; dashed = 0.2 represent the max-normalized levels. The dashed-line box highlights the mobile LHCII1,2 (Lhcb1.3/1.4/2.4) protein subunits. The solid-line box highlights bands 7 and 8 (753 and 720 kDa). Supplemental Fig. S1 provides an independent IgG check of the Lhca and Lhcb proteins.

**Figure 2B.**
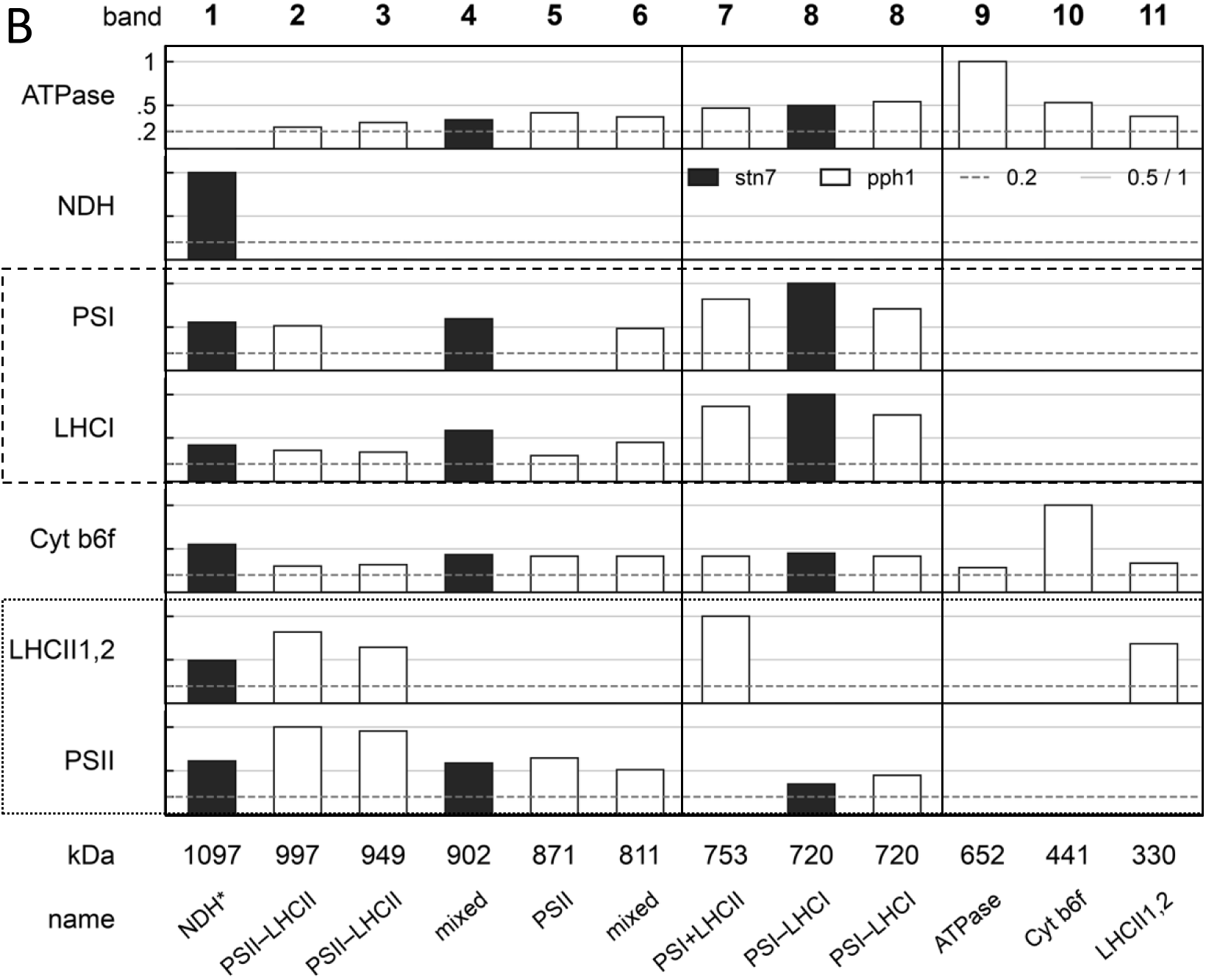
Identification of protein complexes from stroma lamellae by LC-MS/MS after native 1D LDS-PAGE separation. LN-PAGE separated stroma lamellae protein complexes. Fluorescent bands labeled 1 to 11 were analyzed by MS (Methods). Bands were isolated from either the pph1-3 mutant (white) or the stn7 mutant (black). A bar is presented only if the catalytic core of the protein complex is present in the MS dataset. Each row shows the max-normalized peptide counts of proteins sorted by sparkline similarity and merged into the protein complex unit (y-axis label). Mass spectrometric data were normalized to the maximum number of unique peptides per protein. We assigned protein subunits to protein complexes by sparkline correlation (x-axis label), and merged proteins based on literature and crystal structure datasets. We used crystal structures of PSII (PDB entry 3ARC), PSI (PDB entry 2WSC), and the Cyt b6f complex (PDB entry 4H44) to generate master sparklines representing the mobility pattern of the protein complexes across the molecular-weight range of the native gel. We assigned protein subunits of the NDH complex (Suorsa et al., 2009) and ATP synthase (Binder et al., 1978; Zybailov et al., 2008) based on proteomic studies. Gel-band numbers are shown at the top; band kDa values are shown at the bottom. The graph shows solid grey lines at 0.5 and 1 and a dashed cut-off line at 0.2. The solid-line boxed area over columns represents bands 7 and 8; dashed-line boxes over rows show PSI or LHCI distribution; the dotted-line box over rows shows PSII or mobile LHCII1,2 distribution. The star in NDH* indicates that band 1 shows PSI, PSII, and Cyt b6f complexes in addition to NDH (see Discussion).

To characterize the protein bands, we analyzed protein distribution by gel-blot and, especially, by MS. Immunoblots of the stroma lanes identified the subunits that were resident in the specific bands (Fig. 1B, Fig. S1). We used phosphothreonine antibodies to detect the class of phosphorylated proteins and showed that CP43, D1, and D2 were phosphorylated in bands 2 and 3 of both genotypes, and that phosphorylated LHCII was present in *pph1* band 7, but was essentially absent in *stn7*. Antibodies directed against the PSI subunit PsaA were found at bands 7 and 8 in *pph1* and mainly at band 8 in *stn7*, with further signal in *stn7* bands 1 and 4. The antibody against the cytochrome *b*_6_ protein subunit of the Cyt b6f complex was strongest at band 10, and the blot showed no comparable signal above 500 kDa. In antibody screens against light-harvesting proteins of PSI and PSII, protein subunits Lhca1–3 followed bands 7 and 8 and were also detected in bands 1–4, but not in the free LHCII at band 11. Signals for Lhcb1 and Lhcb2 were strong in *pph1* at bands 2, 3, 7, and 11 and essentially absent from band 8. In the stn7 mutant, we detected Lhcb1 and 2 proteins in bands 2, 4, and 11 (Fig. S1). We found that antibodies against Lhcb3–5 reacted most strongly with bands 2 and 3. Based on these gel-blot reactions, we labeled fluorescent bands as photosystem I (I), photosystem II (II), photosystem I with LHCII (IL2), and the cytochrome *b*_6_*f* dimer (Y) (Fig. 1B).

### 2.2 Identification of the LHC and core proteins that occupy the bands after LN-PAGE

Mass spectrometry of the excised stroma bands was used because one antibody does not resolve the composition of the protein complex in a gel slice (Fig. 2). Unique peptide counts from those gel slices were analyzed in two ways (Methods). In the supervised reading, the unique peptide counts per gel slice were arranged in rows, and counts were scaled to their own maximum. A sparkline pattern was generated per row, and the subunits of one complex were summed, which showed an equal sparkline and, in addition, resembled the catalytic pair or a structural subunit of a complex deposited in the PDB database (Fig. 2B). In the unsupervised reading, the same counts were mathematically clustered by the shape of their profiles across the bands, without reference to the subunit list of a databank-deposited protein complex (Table S1, Fig. S2).

We first separated all light-harvesting proteins to differentiate the distribution of mobile Lhcb proteins in the membrane complexes. In the stroma membrane dataset, we scaled each Lhc gene to its own maximum, and bands 1–11 fall into three groups (Fig. 2A). Lhca1, Lhca2, Lhca3, and Lhca4 peak at band 8 and remain high at band 7 in *pph1*. Lhca5 reaches its own maximum in band 1 in *stn7* (four unique peptides), is lower in *stn7* band 4 and in both cuts of band 8, and is absent from band 7 in *pph1*. This count is below the clustered-tree threshold, so Lhca5 does not appear in Table S1. Lhcb1.3, Lhcb1.4, and Lhcb2.4 are at their stroma maximum in *pph1* band 7, are high in bands 2 and 3, and at band 8 remain between 0.13 and 0.26 of that maximum. They are also above the 0.2 line of the stroma scale at *stn7* band 4. The MS analysis of the Lhcb1,2 series differentiates well between the unique PSI pair in bands 7 and 8 and the higher-order assembly states of PSI and PSII complexes in bands 1-4. In addition, Lhcb3, Lhcb5, and Lhcb6 are highest in bands 2 and 3 and are low in bands 7 and 8. The three Lhcb4 gene products and Lhcb5 and 6 also do not follow the band-7 maximum of Lhcb1 and 2. Except for Lhcb4.3, which is low in band 2, all Lhcb4-6 proteins are high or second-highest in bands 2 and 3 and remain above half that maximum in *stn7* bands 1 and 4. Lhcb4.3 is highest in band 3 and is absent from bands 7 and 8. The grana series was recorded for *pph1*. Its tree contains no 753/720 kDa PSI pair, and the LHCII1,2 bar is Lhcb1.3 and Lhcb2.4 (Fig. S2). We did not search the mass spectra for phosphorylation sites, but we performed a phosphothreonine analysis by gel blot after LN-PAGE (Fig. 1B). We found that in *pph1* band 7 carries phosphorylated LHCII, but despite the differentiation of the Lhca1-3, and Lhcb1-6 proteins in Lhca and Lhcb-specific immunoblots, and identification of Lhc proteins by MS (Fig. 2A), we did not differentiate which of the subunits carries the phosphate (Figs. 1B, Fig. S1).

The supervised assembly of the rows was then applied to assemble the core of the protein complexes in each of the bands (Fig. 2B). The complexes were assembled when the core proteins were present. PsaA and PsaB for PSI, psbA-C for PSII, PetA and PetB for cytochrome *b*_6_*f*, AtpA and AtpB for ATP synthase. LHCI, NDH, and LHCII1,2 assembled into a master complex when at least two of their subunits each reached 0.2 of that protein’s maximum per gel bands 1–23. On that scale, PSI and LHCI occupy bands 7 and 8 together and are also present in bands 1, 2, 4, and 6. LHCI was also scored in bands 3 and 5, where PsaA and PsaB do not both pass the bar. PSII is highest in bands 2 and 3, is absent from band 7, but is still detected in both cuts of band 8. The LHCII1,2 row identifies its presence in bands 1, 2, 3, 7, and 11. It is not found in band 4 or band 8, and on the full-gel scale, fewer than two of Lhcb1.3, Lhcb1.4 and Lhcb2.4 reach 0.2 of a maximum that sits in the grana, even though the stroma-scaled gene plot still shows all three at band 4 (Fig. 2A).

NDH is confined to band 1, and band 1 also contains PSI, PSII, and cytochrome *b*_6_*f*. Cytochrome *b*_6_*f* is scored in every stroma band. Its maximum is in band 10, the complex’s dimeric form. The high-mass bands lie between 0.30 and 0.55 of that maximum. For the Cyt b6f complex, *stn7* band 1 is the highest, at 0.55, and band 4 is at 0.43. The two cuts of band 8 are alike, at 0.45 in *stn7* and 0.42 in *pph1*. The data also show that when more than one row is filled, the band represents a basket of complexes that share similar in-gel migration. The name we assign to each band is the majority reading of that basket (Fig. 2B).

In the unsupervised assembly, the clustered tree uses the same stroma counts and keeps a protein only when its maximum in the window is at least 10 unique peptides (Table S1). The distance given for a complex is the height of the bar drawn in Fig. S2. Every leaf under that bar is inside the distance. A higher join is a larger set, and its height is given with the leaves that enter there.

The LHCII1,2 bar is Lhcb1.4 and Lhcb2.4 at *d* = 0.19. Lhcb1.3 joins that pair at *d* = 0.37. All three are maximal in band 7 and remain high in band 11, and band 8 stays between 0.13 and 0.26 of that maximum (Table S1). The PSII bar is drawn at *d* = 0.61. It contains D1, D2, CP43, CP47, PsbO-1, PsbO-2, CP26, CP24, Lhcb4.1, Lhcb4.2 and TLP18.3, and it occupies bands 2 and 3. Lhcb3, PsbS and ascorbate peroxidase 4 join this neighborhood at *d* = 0.91 and lie outside the bar. Lhcb4.3 lies outside both nodes. The PSI–LHCI bar at *d* = 0.56 contains PsaA, PsaB, PsaC, PsaD-1, PsaF, PsaH-2, PsaL, PsaN and Lhca1–4. PsaE-1 and PsaE-2 lie outside it. The complex occupies bands 7 and 8. The NDH bar is drawn at *d* = 0.11 over NdhH, NdhM, Ndf2, NDH48, PPL2 and NdhU (AT5G21430), and these leaves occupy band 1. NdhU, also called CRRL, is a J-domain subunit of the NDH electron-donor subcomplex (Yamamoto et al., 2011; Ifuku et al., 2011). The cytochrome *b*_6_*f* bar at *d* = 0.71 contains PetA, PetB, PetC and Psb33 (AT1G71500). Psb33 carries a Rieske domain and is not the cytochrome subunit PetC. Tic62 (AT3G18890) joins them only at *d* = 0.92. The cytochrome maximum is at band 10, and this bar is separate from the PSI–LHCI bar. The ATP synthase bar is AtpA and AtpB at *d* = 0.17, in band 9. AtpD and AtpE join that pair at *d* = 0.34, AtpC joins at *d* = 0.47, and F0 B′ (AT4G32260) and AtpF join at *d* = 0.70. The distance 0.70 is the full subunit set. The bar drawn for ATP synthase remains AtpA with AtpB. The protein leaves FNR, FtsH, VAR1, AOS, and HCF136 form a separate group that occupies mainly bands 9–11, and that group is not named as a photosystem complex. Bands 1, 4, and one cut of band 8 resemble the *stn7* slices in this tree; the other stroma band analysis is based on *pph1* data. The antibody column presented in Table S1 shows the identification of protein subunits in the gel blots (Fig. 1B and Fig. S1).

The supervised rows and the unsupervised tree do not return the same grouping of the peptide counts. In the supervised analysis, the cytochrome *b*_6_*f* protein subunits are localized to reside in the high-mass bands, while the tree groups the protein leaves outside PSI and keeps PSII visible at band 8. The tree separates Lhcb1.3, Lhcb1.4, and Lhcb2.4 from Lhcb3, CP26, and CP24, which the PSII row of Fig. 2B presented in one trace, and the unsupervised analysis leaves the low-mass FNR–FtsH group without a photosystem name. The sparkline reports where a published complex sits. The tree shows whether two complexes share one profile. The Discussion sets out where the two readings differ and which difference identifies a particle.

### 2.3 Separation and spectral analysis of Chl protein complexes by liquid iZE-FFE

Analysis of band composition, based on both core and LHC subunit distribution, indicated that PSI and PSII complexes co-separated in the same molecular-weight bands, and that the distribution of the Cyt b6f complex was especially unclear. We therefore applied the Digitonin supernatants of *pph1* and *stn7* to charge-only separation using interval-zone free-flow electrophoresis (iZ-FFE). The buffered liquids were collected in a microtiter plate in fractions of 200 to 300 µl, with fraction 1 placed at the anode side of the separation chamber and fraction 96 placed at the cathode side of the chamber. The pH of the fractions resembles the charge axis for the separation of the protein complexes, with a positive sign at the anode and a negative sign at the cathode, and the charge axis is used for analysis of the later free-flow plots (Fig. 3A). The spectral absorbance of the iZ-FFE fractions was recorded at 440 nm and resembled the visible bench record for the plate.

**Figure 3A.**
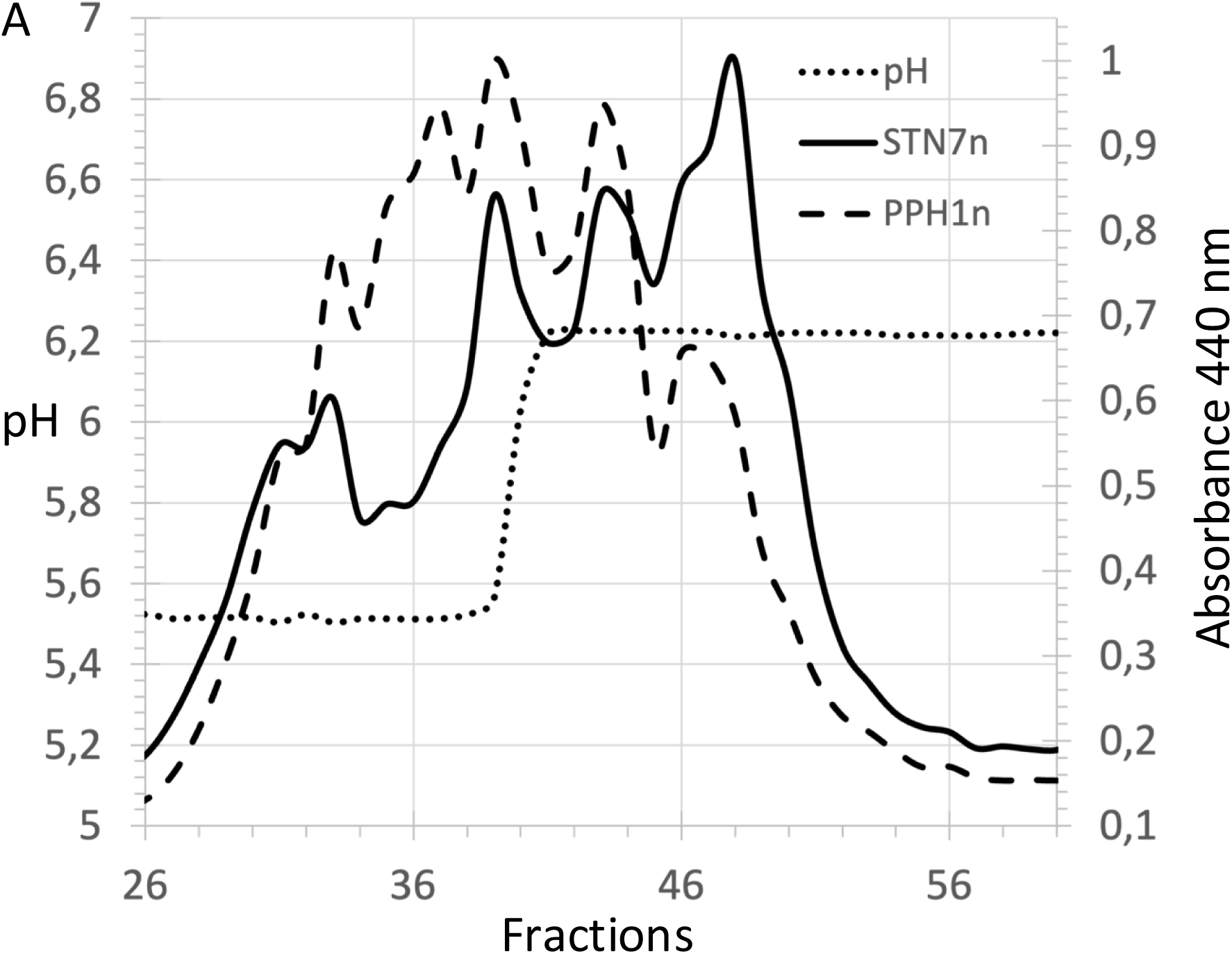
1D iZE-FFE separation of a 16 mM Digitonin thylakoid membrane extract. The stroma lamellae of thylakoid membranes (400 µg Chl) from Arabidopsis mutants stn7 and pph1 were extracted from membranes using 16 mM Digitonin (methods). A volume of 100 µl corresponding to 55 µg Chl was separated by iZE-FFE. The pH and absorbance at A440 nm were recorded. The separation medium was collected (300 µl fractions) in a 96-well MTP. The absorbance of fractions 26 to 60 was plotted after spectra max.-normalization (STN7n, PPH1n).

From the fluorescence emission spectra recorded at 77 K, we extracted the 681 nm (LHCII) and 731 nm (LHCI) regions from each spectrum and plotted them separately along the fraction number/charge axis (Fig. 3B). Absorbance spectra at 440 nm reflect the distribution of Chl-binding protein complexes along the iZ-FFE charge axis and nine absorbance peaks were assigned to the complexes separated from pph1 and stn7 stromal membrane extracts (Fig. 3A). The peaks between the anodic fractions 32 - 44 with a maximum in fraction 37 were increased in pph1, whereas the absorbance peaks at the cathodic side between fractions 44-60 with a maximum in fraction 49 were increased in stn7 (Fig. 3A). Comparison with the pH profile across the charge axis showed that the anodic peak in the pph1 to stn7 difference was localized on the pH 5.5 plateau, whereas the peak in the stn7 to pph1 difference was localized on the pH 6.2 plateau (Fig. 3A). The increased anodic mobility in pph1 indicated that some pph1 complexes were more negatively charged. Fluorescence spectroscopy at 77K was then applied because it allowed monitoring of the PSI distribution via the LHCI fluorescence at 731 nm and the LHCII fluorescence at 681 nm (Fig. 3B). The 2D heatmap of fraction/charge axis over wavelength clearly corroborated the increased anodic shift, and especially the LHCI signal distribution at 731 nm showed that three complexes related to PSI in fractions 36, 38, and 44 increased in pph1 relative to stn7, and the fluorescence in fraction 49 increased in stn7 relative to pph1 (Fig. 3B). This finding indicated three protein bands with increased fluorescence in pph1, while in the LN-PAGE analysis, only band 7 showed increased fluorescence emission in pph1. We therefore identified the protein complex composition of the iz-FFE fractions.

**Figure 3B.**
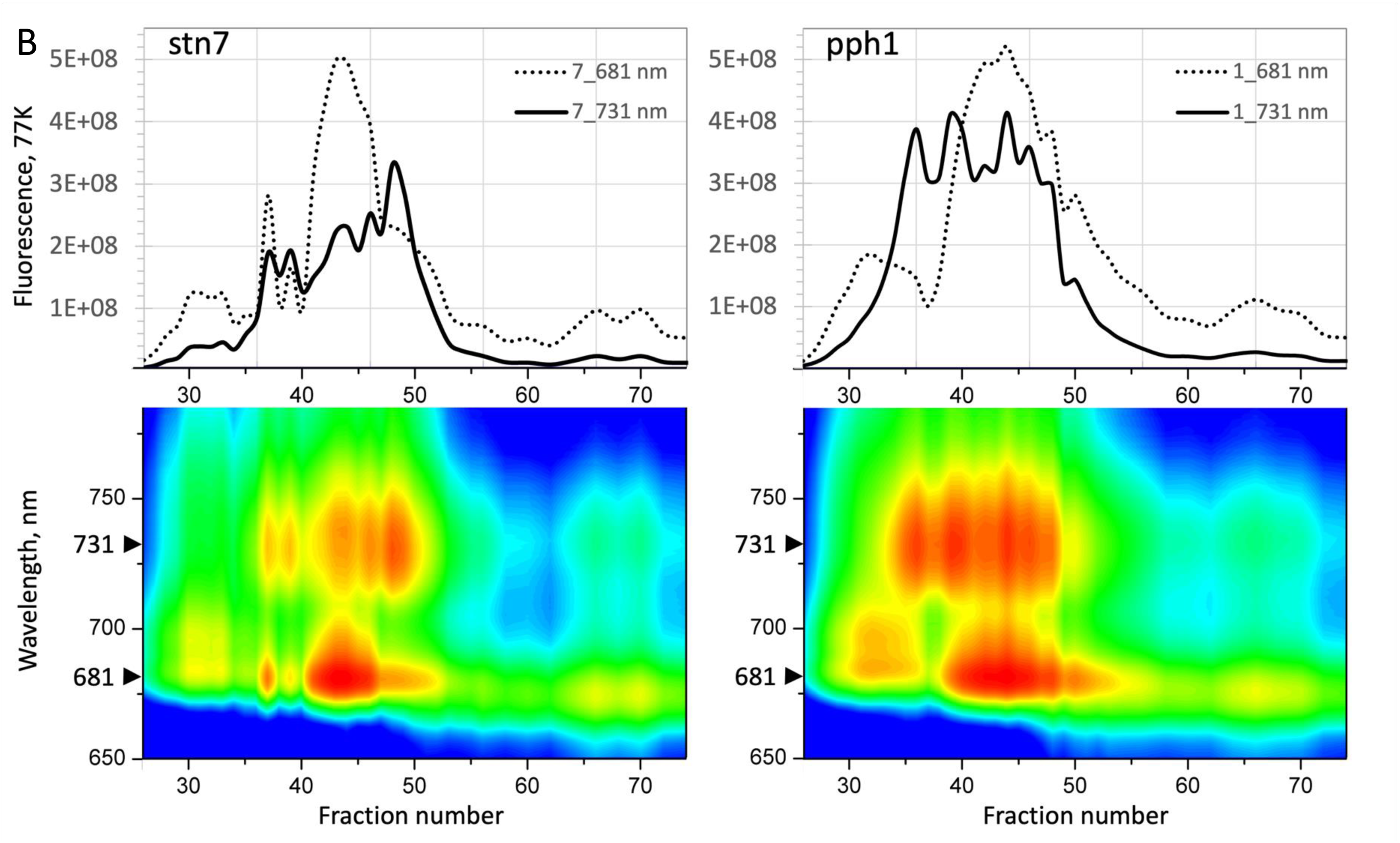
77 K emission of Chl protein complexes from stn7 and pph1 after 1D iZE-FFE. The protein complexes of stn7 and pph1 (Absorbance, Fig. 3A) were separated by iz-FFE (fraction number: 26 to 74) and each fraction scanned for 77K fluorescence emission (Wavelength, 650-800 nm) upon excitation at 440 nm (bottom panels). Two emission spectra (Fluorescence, 77K) were extracted at wavelengths (nm) of F681 = LHCII/PSII and F731 = PSI–LHCI and plotted separately (top panels, stn7, pph1) on the same fraction number basis.

### 2.4 MS determination of protein complexes co-migration in iZE-FFE

Two iZ-FFE separations of the solubilized stroma membranes of pph1 and stn7 were conducted, and the even fractions in the range from fraction numbers 30–60 were digested in solution and quantified by label-free mass spectrometry. Here, we used fmol-like units rather than the unique-peptide count method applied to the gel slices. In the supervised analysis of the dataset, the same protocol was followed, and the fmol units were grouped from the same subunit lists as shown (Fig. 2B). An identified protein in a fraction was scored only when the catalytic core of the protein complex was identified as well, and the rows of proteins constituting the protein complex were scaled to their own maximum (Fig. 4). The normalized distribution quantity of each protein complex identified per fraction in *stn7* (Fig. 4, solid line) and *pph1* (Fig. 4, dashed line) was plotted together to directly compare the distribution changes along the fractions/charge axis (Fig. 4). PSII and LHCII1,2 of pph1 and stn7 occupied an anodic region between wells 30 and 44 with stn7 extending the distribution of both protein complexes further towards the anode. In contrast to PSII, LHCII1,2 showed a bimodal distribution with a second peak in the cathodic region between fractions 48 and 56, and maxima at 50 (stn7) and 52 (pph1). This second LHCII1,2 mode showed a parallel mode in the same charge region in PSI and LHCI for the pph1 complexes. Three protein complexes, PSI and LHCI, as well as the Cytochrome *b*_6_*f*, showed an overall increase in quantity throughout the investigated separation space between fractions 36 and 60. PSI and LHCI showed the most similar charge-dependent distribution, followed closely by the Cyt b6f complex, which showed a distribution similar to the PSI-LHCI distribution in both pph1 and stn7. The ATPase and NDH complexes peaked at fractions 40 and 46, respectively, and the distribution pattern was most similar between pph1 and stn7 in ATPase. Interestingly, we noticed that the order of complex mobility along the anode-to-cathode axis appeared reversed relative to the order in which the complexes separated in LN-PAGE. While PSII showed the highest mobility toward the anode, the ATPase, NDH, and LHCII1,2 complexes showed progressively slower mobility, while PSI and LHCI populated a large charge volume toward the cathode, clearly distinguishing the PSI-related complexes from PSII and ATPase. The analysis indicated a symmetric distribution for cathodic LHC1,2 and the more anodic ATPase and NDH distributions, with about +/- 3 fractions considered overlapping relative to the peak.

**Figure 4.**
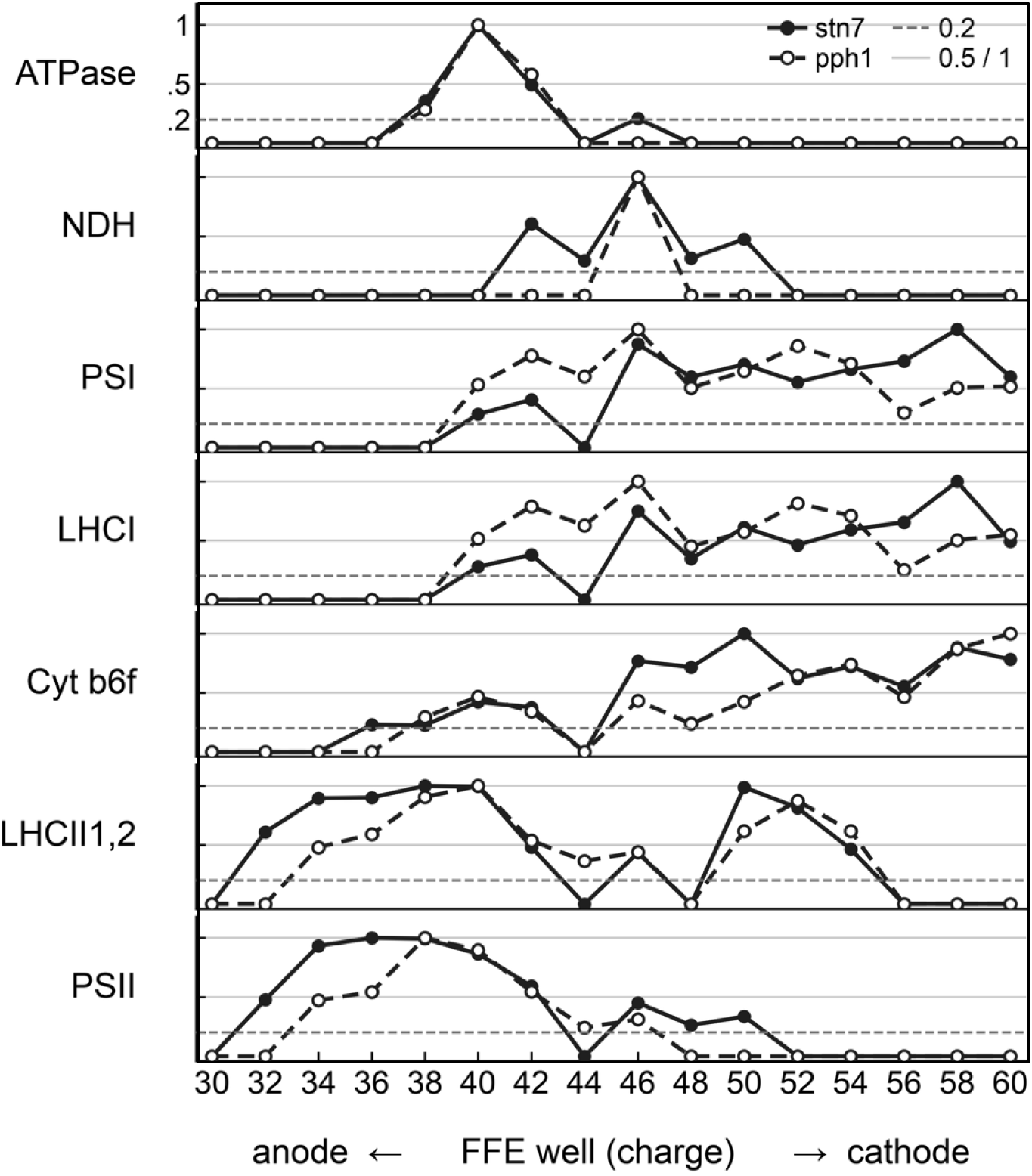
Charge distribution of protein complexes after 1D iZE-FFE fractionation. A microtiter plate volume (300 µl) of even wells 30–60 was concentrated on a Vivaspin500 (10K), and proteins were processed for MS identification using a FASP protocol (methods). Proteins were identified by peptide sequencing and quantified using label-free peptide intensities. Proteins were merged into protein complexes ATPase, NDH, PSI, LHCI, Cyt b6f, PSII, and LHCII1,2 using the protein database deposited structural data (pdb.org). The sum of all protein counts per complex was max-normalized per FFE fraction. Counts from stn7 are shown as a solid line with filled data marks; counts from pph1 are shown as a dashed line with open data marks. Relative count levels are indicated at 0.5 and 1 (grey lines), and a 20% cut-off level is shown (dashed line at 0.2). The FFE fractions are shown with increasing numbers from the Anode (left) to the Cathode (right).

In the unsupervised analysis, we called the tree and applied hierarchical clustering to the same dataset. We set the cut-off to 2 fmol-like units (Fig. S3A) and collapsed each majority bar to the mean of its member traces, rescaled it to its own maximum, and plotted it on the same axis of fractions/charge. We present the leaf order, distances, and the uncollapsed heatmaps (Fig. S3B and Fig. S3C). The tree returns the same complexes as the supervised rows, with two restraints that the supervised rows do not make (Fig. S3A, Fig. S3B and Fig. S3C). The first is the minor antenna. The supervised row averaged Lhcb3, the four CP29 accessions, CP26 and CP24, because those charge profiles correlated (*r* ≥ 0.94 for CP24 against CP29 and Lhcb3) and lacked the cathodic rise of Lhcb1 and Lhcb2. On the tree, the short node was narrower. It contained two CP29 accessions and Lhcb3, at *d* = 0.33 in *stn7* and at *d* = 0.26 in *pph1*. CP24 and CP26 are positioned outside that node. The PSII bar is the reaction center with PsbE (D1, D2, CP43 and CP47; *d* = 0.17 in *stn7* and *d* = 0.16 in *pph1*). The minor-antenna node is falling outside that bar. Both occupy the anodic wells, which is why the oxygen-evolving subunits and this anodic LHCII pool overlap on the fraction axis. One CP29 accession sits nearer the PSII reaction center on the tree than its charge profile does (Fig. S3B and Fig. S3C).

The second restraint was the FNR-like protein 2. LHCII1,2 on the tree was found bimodal, with an anodic maximum and a second rise near wells 50–52. In *stn7* the three Lhcb1 and Lhcb2 leaves joined at *d* = 0.73, and the Lhcb1 plus Lhcb2 pair alone at *d* = 0.64. CB1.1 peaks at well 50, while CB1B1 and Lhcb2 peaked at wells 38–40 and still rose near well 50. That is one clade. In *pph1*, two leaves joined at *d* = 0.44, and the next join of that neighborhood included FNRL2 (Fig. S3A). FNRL2 kept the cathodic shoulder and correlated with Lhcb2 in *pph1* (*r* = 0.94). FNRL1 stayed anodic. The mean of wells 48–54 was 0.09 in *stn7* and 0 in *pph1*. Cytochrome b_6_f occupied its own bar on this charge tree in both genotypes (*d* = 0.20 in *stn7* and *d* = 0.38 in *pph1*). Neither the supervised rows nor this tree placed PSI–LHCI, the cathodic LHCII1,2 population, and cytochrome *b*_6_*f* in one node.

The same trees placed several leaves beside those bars (Fig. S3B and Fig. S3C). The PSI bar peaked at well 58 in *stn7* and at well 46 in *pph1*. The NDH bar peaked at well 46 in both genotypes, so the *pph1* PSI maximum fell in the NDH well, while the *stn7* PSI maximum was localized to the cathode side of it. The CURT1A protein (AT4G01150; Armbruster et al., 2013) was found outside the PSI bar in both genotypes. Its profile correlated with that bar (*r* = 0.70 in *stn7*; *r* = 0.86 in *pph1*). In *pph1* it peaked at well 46 and was the leaf beside PsaG. In *stn7* its maximum was at well 42, and the cathodic side of the profile remained high.

AT5G02160 was found outside the ATPase bar in both genotypes. It peaked at well 40 in *stn7* and at well 38 in *pph1*, in the anodic window that also contained PSII, the minor antenna and the FtsH proteins, and in *pph1* the leaf was placed beside PsbO-1. Proteins that peak in this narrow anodic zone correlated with more than one bar, so the correlation left AT5G02160 outside ATP synthase and outside PSII. Rhodanese was present only on the *stn7* tree. It lay immediately before F0 subunit B and outside the ATPase bar, which was the five-subunit node at *d* = 0.14 (AtpA, AtpB, AtpD, AtpE and AtpF). The *stn7* NDH bar (*d* = 1.06) also covered NdhU and the lumenal NDH subunits PnsL5 (CYP20-2; AT5G13120, leaf label F4K2G0) and PnsL4 (FKBP16-2; AT4G39710, leaf label F4JW56) (Ifuku et al., 2011). The same bar covered HCF136, a PSII assembly factor, which is not an NDH subunit.

The spectroscopic plate, the liquid mass spectrometry, and the free-flow separation loaded on the second gel were prepared on different days, from separate membrane isolations and detergent extracts. The fraction numbers are the run numbers. We aligned them by two correspondences. Liquid-MS well 48 corresponded to well 41 of the second-dimension gel, a shift of seven wells. Spectroscopic fractions 36 and 38 corresponded to wells 38 and 40 of that gel, a shift of two wells. The two shifts agree. Spectroscopic fraction N fell on liquid-MS fraction N + 9 and on second dimension well N + 2.

Under that alignment, the *stn7* fluorescence increased at spectroscopic fraction 49 fell on liquid-MS well 58, where PSI and LHCI of *stn7* were at their maximum and the *pph1* values were about half (Fig. 4). The three *pph1* fluorescence increases, at spectroscopic fractions 36, 38 and 44, were falling on liquid-MS fractions 45, 47 and 53. Fraction 45 was beside well 44, where PSI and LHCI were present in *pph1* and absent in *stn7*. Fraction 53 was localized beside well 52, where *pph1* PSI and LHCI remained above the *stn7* traces and where *pph1* LHCII1,2 had its cathodic maximum. Fraction 47, between wells 46 and 48, is where PSI was high in both genotypes. The liquid MS recorded only even-numbered wells, so a difference peak placed on an odd fraction was not resolved as its own particle, and the three *pph1* peaks were not three separate PSI complexes in that matrix. On the second-dimension gel, the same alignment placed spectroscopic fractions 36 and 38 on wells 38 and 40, anodic of the slices that were cut for mass spectrometry, spectroscopic fraction 44 on well 46, at the maximum of the *pph1* 753 kDa series, and spectroscopic fraction 49 on well 51, where PSI was high in both genotypes (Fig. 5A and Fig. 5B).

**Figure 5A.**
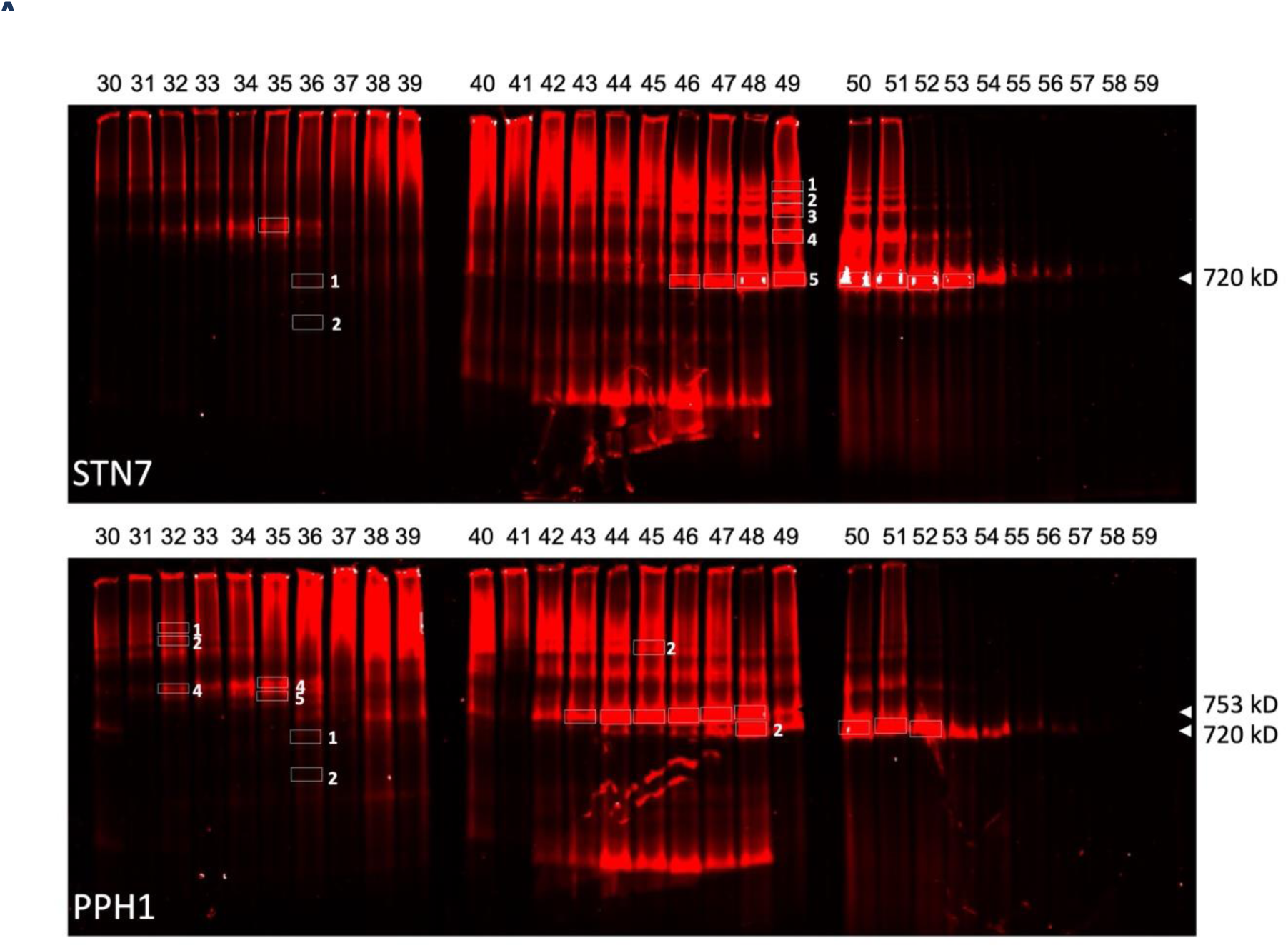
LN-PAGE of iZE-FFE fractions from mutants stn7 and pph1. A volume of 200 microliters was retrieved per FFE fraction 30 to 60, concentrated 8-fold to 25 microliters, and protein complexes were processed for separation by LN-PAGE overnight (methods, wells 30 – 59 shown). Gels were scanned using the LiCOR CLX scanner (Ex 680, Em > 700 nm). Proteins selected for MS sequencing analysis are boxed. A horizontal series of protein complexes was selected for MS identification of the protein composition at 720 kDa (stn7, lanes 46 – 53); a series of five bands with increasing molecular weight in lane 49 (49-1 to 49-5); and bands in lanes 35 and 36-1 and -2 (stn7, solid line boxes). For pph1, we selected bands 43–48 at 753 kDa and bands 48-2–52 at 720 kDa. In addition, individual protein bands were selected for MS analysis from the bands in lanes 32-1,2, and 4, 35-4, and -5, 36-1, and -2, and 45-2 (pph1, solid line boxes). Each well is a charge window from the anode (left) to the cathode (right).

**Figure 5B.**
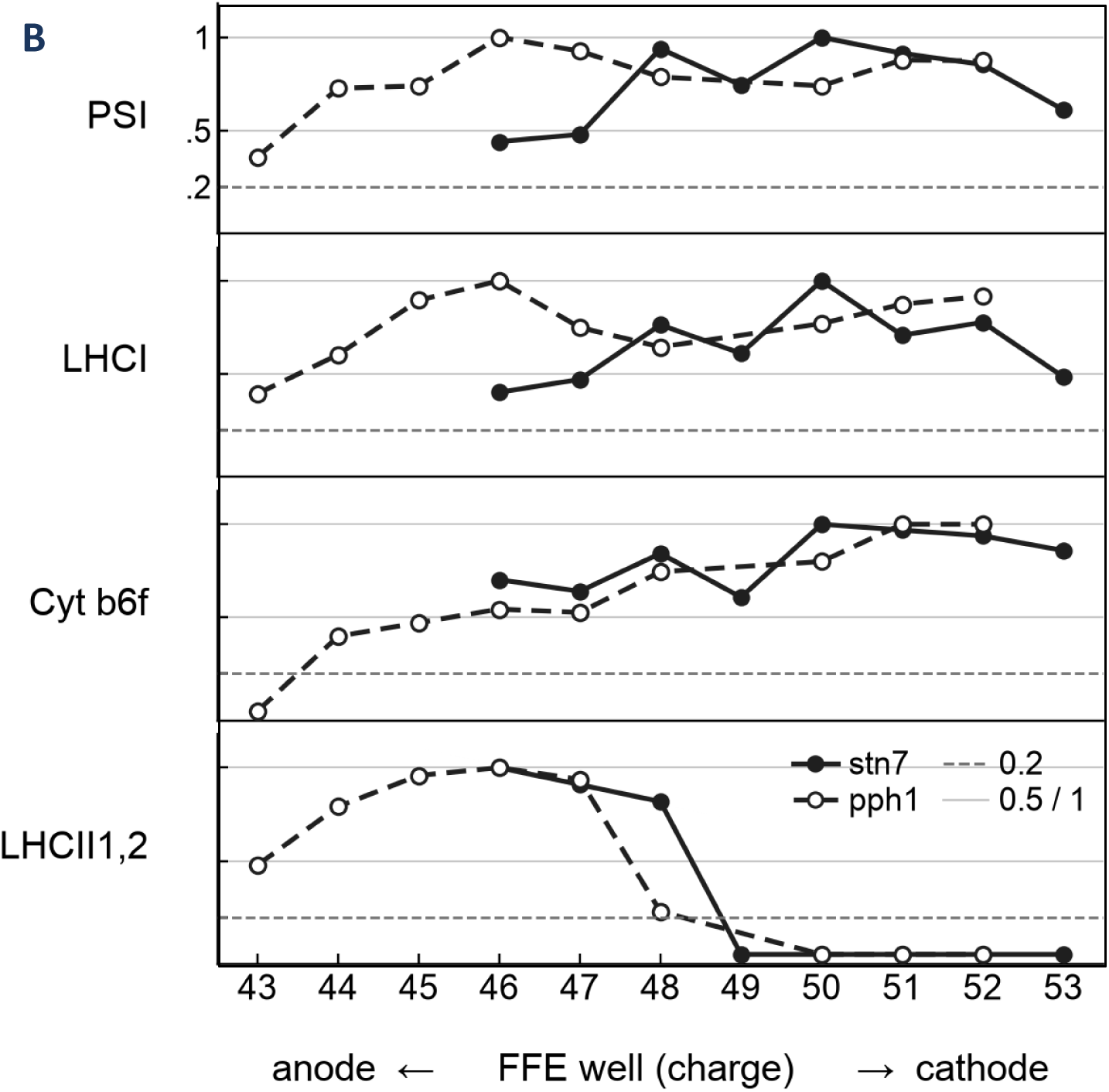
Charge distribution of protein complexes after 2D iZE-FFE/LN-PAGE fractionation. The gel slices were cut after Coomassie staining (gel regions shown as white boxes in fluorescent Fig. 5A and in Coomassie-stained Fig. S4). Gel slices were processed by in-gel digestion, and peptides were subjected to MS sequencing analysis (methods). Proteins identified by peptide sequencing were quantified using spectral counts and merged into the protein complexes PSI, LHCI, Cyt b6f, LHCII1,2 on the basis of published protein complex composition. The sum of all peptide counts per complex was max-normalized per fraction. Counts from stn7 are shown as a solid line with filled data points, and counts from pph1 as a dashed line with open data points. Relative count levels are indicated at 0.5 and 1 (grey lines), and a 20% cut-off level is shown (dashed line at 0.2). The FFE fractions increase from the Anode (left) to the Cathode (right).

We tested this question by subfractionating the iZ-FFE fractions into distinct molecular-weight subclasses using a second-dimension LN-PAGE approach. We reasoned that an interaction between the lower-molecular-weight LHCII or Cyt b6f complex and the larger PSI-LHCI complex should be detectable as a molecular-weight shift in LN-PAGE.

### 2.5 The 720 kDa class is PSI–LHCI–cytochrome *b*_6_*f* in both genotypes, and the 753 kDa class binds LHCII1,2 in *pph1*

In the first preparatory step for LN-PAGE separation, we concentrated free-flow volumes about 8-fold using 100K cut-off filter membranes. After separating the complexes, we first scanned the native gel for Chl fluorescence (Fig. 5A), then stained the gel proteins with colloidal Coomassie (Fig. S4). We cut selected Coomassie band slices from the gel for in-gel digestion and MS analysis, especially with a focus on investigating the PSI complex interactions (Fig. 5B). In the analysis of the distribution of fluorescence in the native gels, a series of bands at 720 kDa was dominant in the stn7 mutant in the cathodic region of the FFE axis between fractions 46 to 54 (Fig. 5A). A similar series of bands was present between fraction 47 to 54 and in the same molecular weight region at 720 kDa in pph1. In addition, a series of bands was dominant in fractions 42 to 48 with a molecular weight of about 753 kDa in pph1 (Fig. 5A). Interestingly, the fractions containing the 753 kDa band were shifted towards the anode. This finding established a direct link between band 7 (753 kDa) and band 8 (720 kDa) in the 1D LN-PAGE approach in pph1, and between band 8 (720 kDa) in stn7 (Fig. 1A and B). With the identification of the PSI-LHCI-LHCII1,2 complex in band 7, the finding of an anodic shift in the 2D approach directly indicated that the iZ-FFE separation shifted band 7 relative to band 8 based on the interaction of the LHCII1,2 complex with the PSI-LHCI complex (Fig. 5A). The stn7 mutant also showed a series of strong fluorescent bands in fractions 48 to 51, with increasing molecular weight relative to the 720 kDa base. The origin and nature of this band sequence were unclear. With the 1D LN-PAGE gel clearly defining the band 7 and 8 protein complex content, we wanted to determine the composition of these bands after 2D iZ-FFE LN-PAGE separation.

In our analytical MS approach, we again followed the strategy to investigate the protein distribution using our supervised and unsupervised HCA approach. We focused on two groups: the 720 kDa PSI class in the more cathodic wells between 46 and 53 in stn7, and the cathodic 720 kDa plus more anodic 753 kDa PSI class protein bands in *pph1*. For this analysis, we did not recover the NDH, PSII, and ATP synthase cores that were present in the 2D iZ-FFE separation in the anodic regions between fractions 30 and 44. The remaining rows of proteins that were combined in the analysis are PSI, LHCI, cytochrome *b*_6_*f*, and LHCII1,2, and the analysis of the complexes related to the pph1 band 7/8 (Fig. 5A, dashed line) and to the stn7 bands 8 (Fig. 5A, solid line) were combined in one graph. In *pph1*, the LHCII1,2 complex shows mobility shifted toward the anode relative to the *stn7* LHCII1,2 complex. The graph does not show a shift in the complex’s molecular weight because, based on the fraction separation of bands 7 and 8 in pph1, the LHCII1,2 content is represented in both bands. Furthermore, the distribution of LHCII1,2 in the two mutants is presented on a normalized axis, which shows that relative to stn7, LHCII1,2 in pph1 is shifted to the anode in mobility corroborating the maintained phosphorylation status of the LHCII1,2 complex in the 753 kDa band in fractions 42 to 47 (Fig. 5A). With respect to the Cyt b6f complex, band 7 analysis revealed that in *stn7*, the PSI, LHCI and the cyt *b*_6_*f* complex show a very similar charge profile. Most interstingly, the joint charge profile of the three complexes shift to the anode together in *pph1* plus the profiles of PSI and LHCI show an overlay with the anodic shift of the increased LHCII1,2 level in pph1. This similarity in LHCII1,2 distribution with PSI and LHCI, which we noted in the 1D iZ-FFE analysis (Fig. 4), reappeared here but was now directly visible alongside the anodic displacement of the LHCII1,2 complex (Fig. 5B).

In a secondary attempt to further characterize the PSI-LHCI-Cytb6f complex identified in the 720 kDa bands, we cut five gel slices from the fluorescent ladder in *stn7* well 49. We clustered these slices separately, from 720 kDa slice 49-5 to 49-1 (Fig. S5). We also applied a cut-off floor of 10 counts in this window, and twenty proteins passed it. PsaA and PsaB joined at *d* = 0.28, together with PsaD-1, PsaE-2, and Lhca1. Lhca3 followed in the next shells of that group. PetA (12 to 17 counts) and PetC (5 to 14 counts) were present in every slice. PetB was also present in every slice (5, 2, 3, 2, and 8), but its maximum of 8 stayed below the floor, so this ladder drew no PetA plus PetB bar. These five slices did not belong to one apparent-mass class, and the MS analysis did not resolve the origin of the increased molecular mass. Slice 49-5 is the 720 kDa band, and slices 49-1 to 49-4 were positioned at increased molecular mass. We therefore did not summarize the counts from this ladder of gel slices into a single composition. Lhca2 did not pass the floor level, as it was only two counts, and only in slice 49-5. Lhca4 reached only 9 counts. Lhcb1.2, Lhcb1.5, and Lhcb2.2 stayed at or below 3, so the ladder was not an LHCII series. ATP synthase was a separate clade in the tree at *d* = 0.34 (AtpA, AtpB, and AtpC1). Those subunits were highest in slice 49-1 and were at background level in 49-4 and in 49-5. PsaD-2 was confined to 49-5, at 30 counts, and was absent from 49-1 through 49-4. The four slices above the 720 kDa band therefore carried the PSI and PetA profile of that band, and slice 49-1 adds an ATP synthase profile that the 720 kDa slice lacked.

Unsupervised clustering of the 720 kDa slices showed that PSI–LHCI formed a complex with cytochrome *b*_6_*f* in both mutants (Table 1). Phosphorylated LHCII1,2 binding was a second result, and it was confined to the *pph1* 753 kDa slices. We extracted the *stn7* 720 kDa window from lanes 46–53. The *pph1* 720 kDa window was built from the bands specified in lanes as 48-2 and 50-2–52-2. We measured the 753 kDa window composition from lanes 43–47. In both genotypes, the 720 kDa node is PSI–LHCI–cytochrome *b*_6_*f*. PsaA, PsaB, PetA, and PetB were identified as leaves of that node, together with Lhca1 and Lhca3 (*stn7 d* = 0.50; *pph1 d* = 0.34; Table 1). The proteins Lhcb1 and Lhcb2 were not in this node. In *pph1*, the same node also contained PetC, a P-loop NTPase, and PNDOR. The NTPase and PNDOR co-migrated under the node. Although these two proteins were not considered photosystem subunits, we discuss their potential function. The distances reflected the similarity of the max-normalized count profiles across the window slices. The 753 kDa window was specific to the *pph1* mutant only (lanes 43–47). The tree of that window was also drawn from proteins whose highest count in any one slice was at least 10. Its node contains PsaA, PsaB, Lhca1, Lhca3 and Lhca4 together with Lhcb1.2, Lhcb1.5 and Lhcb2.2, and it also contained PetA and PetC (*d* = 0.38; Table 1). We determined PetB with counts of 5, 6, 6, 6, and 6 across the five gel slices at 753 kDa. The highest value was 6, so PetB did not enter the matrix, and the tree drew no PetA plus PetB bar. The five slices were, however, one apparent-mass class, cut from successive fractions along the charge axis.

**Table 1.**
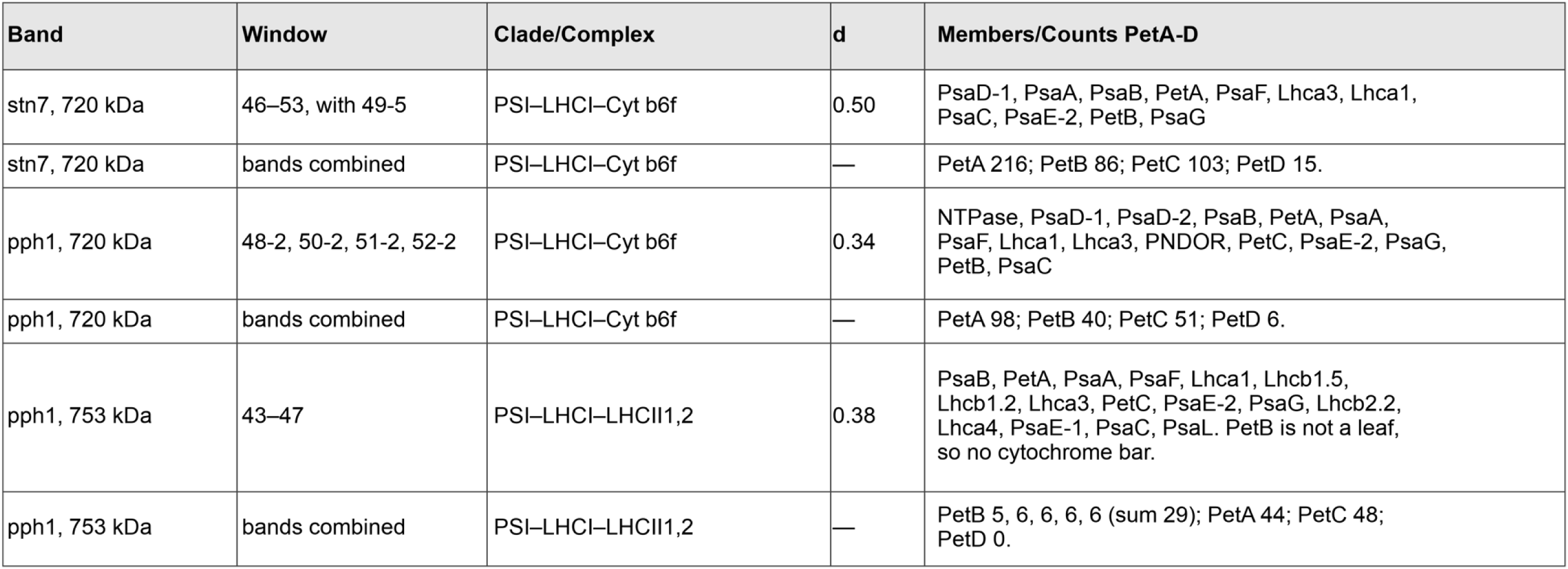
Assignment of the 720 kDa and 753 kDa slices after iZE-FFE and LN-PAGE (Fig. 5C). Each band is entered twice. The tree row lists the slices in Window, the complex in Clade/Complex, the distance in d, and the leaves in Members/Counts PetA-D. The next row has Window “bands combined”, which adds the slices of that one band. Clade/Complex repeats the same complex. Column D is a dash because the row is not a tree node, and Members/Counts PetA-D gives PetA, PetB, PetC, and PetD. The *stn7* 720 kDa window (wells 46–53, with slice 49-5) and the *pph1* 720 kDa window (48-2, 50-2, 51-2, 52-2) are both named PSI–LHCI–cytochrome *b*_6_*f* (*d* = 0.50 and *d* = 0.34). Only *stn7* draws a separate cytochrome bar inside that node. In *pph1* 720 kDa, PetA and PetB already share the node, so no separate cytochrome bar is drawn. NTPase is AT1G77620 and PNDOR is AT1G74470 (CHLP); both are leaves of the *pph1* 720 kDa node. The *pph1* 753 kDa tree (wells 43–47) is PSI–LHCI–LHCII1,2 (*d* = 0.38). PetB is not a leaf of that tree. The two 720 kDa windows are not added together.

For a subunit already assigned to cytochrome *b*_6_*f*, the class composition is the sum of its counts across the slices. PetB was present in all five gel slices (sum 29), while PetA summed to 44 and PetC to 48. PetD contributed no counts in these slices. We therefore concluded that the 720 kDa class is a PSI–LHCI–cytochrome *b*_6_*f* particle in both genotypes. The *pph1* 753 kDa class was the secondary result: the PSI–LHCI bound to phosphorylated LHCII1,2. PetB was present in every slice of that class but the barrier indicating that the gel-shift between the 720 to 753 kDa band to represent the PSI–LHCI–cytochrome *b*_6_*f* particle binding LHCII1,2 was mathematically not written into the tree as petB was not a leaf of the tree.

## 3. Discussion

### 3.1 Room-temperature fluorescence of the gel records chlorophyll, not the complex

Charge separation followed by a second native gel isolates a PSI–LHCI–cytochrome *b*_6_*f* particle from the stroma lamellae of both genotypes, and, only in *pph1*, a PSI–LHCI particle that also binds LHCII1,2. When we scanned the gel for chlorophyll fluorescence at room temperature and, separately, stained it for protein (Fig. 1A), the two images diverged in both directions. Band 22, corresponding to free LHCII in the grana lane, is the strongest fluorescent band in the stroma lamellae, and its protein quantity falls below the SYPRO Ruby signal of the neighboring protein bands. Bands 9 and 10, the ATP synthase and the cytochrome *b*_6_*f* dimer, take up the protein stain and do not fluoresce. A ratio of the two images is therefore undefined wherever one of them is at background, and it is not a chlorophyll-to-protein ratio in the bands where both are visible. Across bands 2, 3, 4, 7, and 8, fluorescence follows the protein stain, the opposite of what would be expected if a chlorophyll-rich, protein-poor antenna were the brightest band. This indicates that fluorescence in the free LHCII band is unquenched and appears to be the only complex in the gel in which excitation-energy moderation by coupling of the LHC proteins to reaction-center core components is absent. Both Bands 7 and 8 showed fluorescence in that scan, and both contain photosystem I (Fig. 1B, Fig. 1C). In a leaf, photosystem I contributes little room-temperature fluorescence, because excitation is trapped by P700. Native gels are nevertheless imaged by chlorophyll fluorescence, including the PSI-containing region (Järvi et al., 2011). The gel shown here reacted to excitation light in the same way. Once the complex is in detergent and in polyacrylamide, the yield can change. Quenching by reaction-center core components may change because some antenna chlorophylls are no longer tightly connected to a reaction-center chlorophyll, or because Chl in contact with the gel can redistribute excitation energy. This study does not separate those contributions. Here, we use the room-temperature fluorescence recording only as a map of chlorophyll-binding material.

The 77 K spectra we recorded from the gel slices are a different measurement. Here, the excised band 7 and band 8 each emit at 680 nm and near 730 nm. At 77 K, the 680 nm band is LHCII emission that is not transferring excitation to a reaction center, and the long-wavelength band is PSI–LHCI emission (Rijgersberg et al., 1979; Grieco et al., 2015). Normalized at 731 nm, band 7 has about 1.9-fold the 680 nm emission of band 8 (Fig. 1C). That comparison is between two excised slices. It does not identify the complex in a room-temperature gel lane, and a 680 nm-rich spectrum of a digitonin extract does not locate the Lhcb protein. The grana tree still contains Lhcb1.3 and Lhcb2.4 (Fig. S2). Low-concentration digitonin releases the stroma lamellae and leaves PSII enriched in the granal residue (Andersson and Anderson, 1980). The genotype difference followed in this paper is the one in that stroma lane. We carried forward bands 7 and 8 because they were characteristic of the mutants, appeared at distinct molecular masses on the gel, and showed a distinct difference in LHCII-specific fluorescence. Band 7 is present in *pph1* and absent from *stn7* and the wild-type stroma (Fig. 1A). Additional bands that are specific to the mutant state are bands 1 and 4 in *stn7*.

### 3.2 A 1D band resembles a basket, while the 2D iZE/LN-PAGE band resolves a protein complex

One crucial finding at the beginning of this work was that digitonin at 16 mM does not return one complex per fluorescent band after LN-PAGE (Järvi et al., 2011). In the MS identification of the protein complexes, we designed the supervised row analysis of the peptide counts to keep a complex visible in every band where its catalytic pair, or at least two of its antenna proteins, passes the threshold (Fig. 2B). That is why cytochrome *b*_6_*f* is scored in every stroma band, from 0.30 to 0.55 of its maximum at the dimer in band 10, and why PSI and LHCI can be drawn as two rows despite the finding that their profiles match. To our surprise, the immunoblot does not show a comparable cytochrome *b*_6_ signal above 500 kDa (Fig. 1B). However, the peptides in the high-mass gel slices are real at a level the blot did not register.

The unsupervised similarity analysis, which builds an HCA tree from the same counts, does not use the PDB subunit list (www.pdb.org) as a constraint (Table S1, Fig. S2). It draws the LHCII1,2 bar on Lhcb1.4 and Lhcb2.4 at *d* = 0.19, and Lhcb1.3 joins at *d* = 0.37. CP26, CP24, Lhcb4.1 and Lhcb4.2 lie inside the PSII bar at *d* = 0.61, and Lhcb3 joins that neighborhood at *d* = 0.91, outside the bar. It groups Lhca1–4 with the PSI subunits and leaves cytochrome *b*_6_*f* outside that PSI group, with the cytochrome maximum still at band 10. It leaves Lhcb4.3 outside the PSII bar, and it leaves the FNR–FtsH group at low mass without assigning a photosystem name according to the similarity definitions set. We concluded that a gel band that shows several filled rows of distinct photosystem complexes represents a basket. The rows indicate which published complexes are present, and the tree shows which proteins share a profile. Neither indicates that we isolated a single complex as a distinct particle on this gel. The same limit applies to blue-native slices cut at 1 mm (Takabayashi et al., 2013).

The two readings disagree in three places, and only the third identifies a particle. On the stroma gel, the sparkline scores cytochrome *b*_6_*f* in the high-mass bands, and the tree leaves it outside PSI, with its maximum at band 10 (Table S1, Fig. S2). On the free-flow axis, both readings keep cytochrome *b*_6_*f* as its own charge class. The tree adds two restraints the sparkline does not: the minor-antenna average used for the supervised row is wider than the clade, which contains two CP29 accessions and Lhcb3, and FNRL2 travels with the *pph1* Lhcb1/Lhcb2 shoulder without being a subunit of that complex (Fig. S3A, Fig. S3B and Fig. S3C). After the second native gel, the sparkline shows that PSI, LHCI, and cytochrome *b*_6_*f* occupy the same 720 kDa wells. The tree shows that PetA and PetB join the PSI–LHCI node in both genotypes (Fig. 5C, Table 1). That join is the identification. The *pph1*-specific 753 kDa class is the second result: phosphorylated LHCII1,2 joins PSI–LHCI, and PetB is not a leaf of that tree. The hierarchical clustering is unsupervised in the sense that the subunit list is applied after the profiles have been grouped. Its value is the refusals on the first two axes and the join on the third. Agreement between the two readings makes the join trustworthy.

**Figure 5C.**
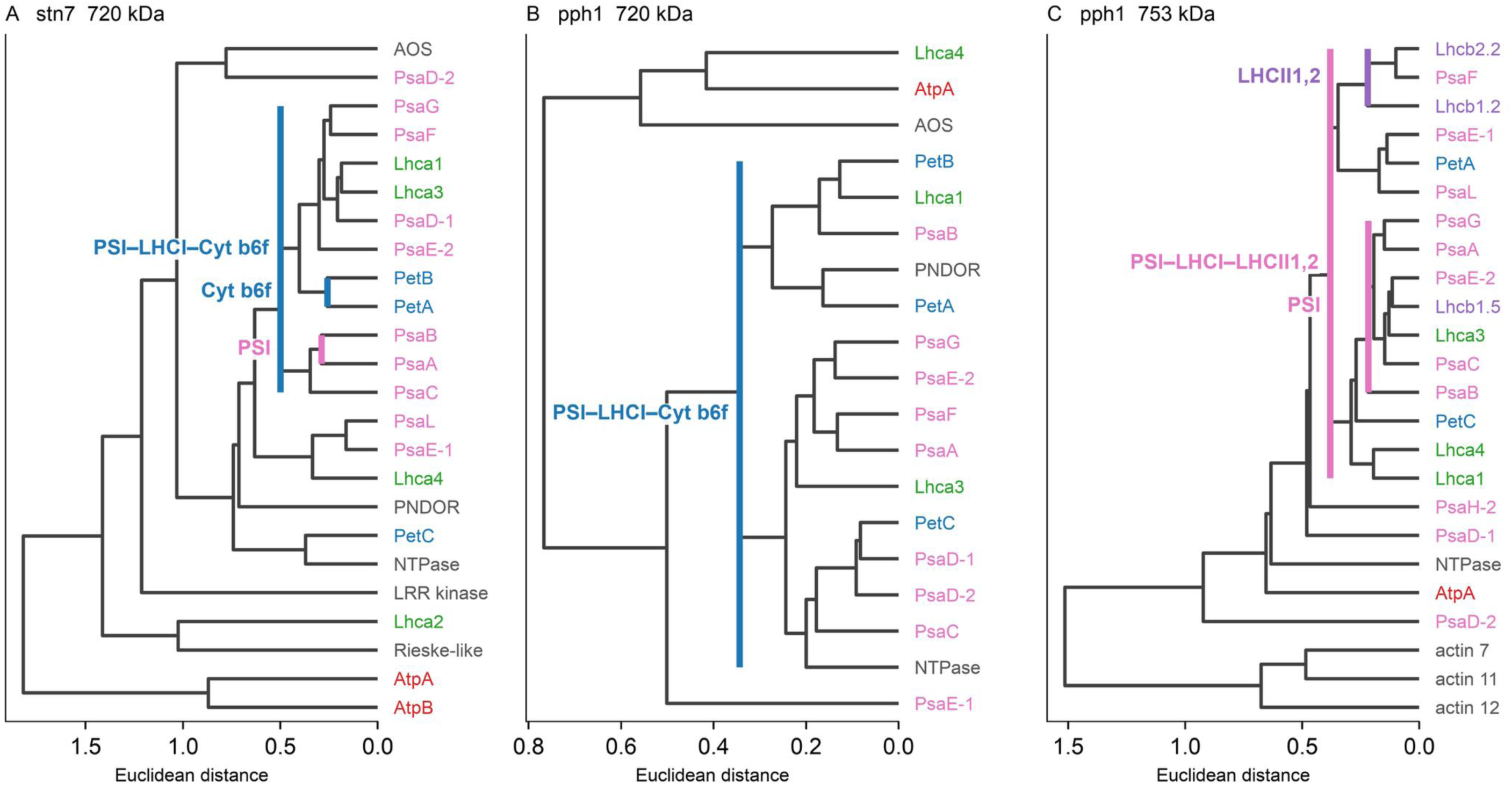
The trees that generate Table 1. Average-linkage Euclidean clustering of max-normalized spectral counts, with a per-slice maximum of at least 10. (A) *stn7* 720 kDa, wells 46–53 including slice 49-5. The outer bar is PSI–LHCI–cytochrome *b*_6_*f* (*d* = 0.50). Inside it, cytochrome *b*_6_*f* is PetA and PetB (*d* = 0.26), and photosystem I is PsaA and PsaB (*d* = 0.29). (B) *pph1* 720 kDa, slices 48-2, 50-2, 51-2, and 52-2. The outer bar has the same name (*d* = 0.34). PetA and PetB already sit in that node, so no separate cytochrome bar is drawn. NTPase (AT1G77620) and PNDOR (AT1G74470, CHLP) are leaves of this node. (C) *pph1* 753 kDa, wells 43–47. The outer bar is PSI–LHCI–LHCII1,2 (*d* = 0.38) and is not a cytochrome bar because PetB is not a leaf. The inner LHCII1,2 bar covers Lhcb1.2, Lhcb2.2, and PsaF. The inner photosystem I bar includes Lhcb1.5. Distance is profile shape. A subunit below the per-slice floor is absent from the tree and can still be counted in the Table 1 sum.

The Cytochrome *b*_6_*f* complex is recovered from both appressed grana membranes and stroma-exposed membranes, unlike the PSII core (Anderson, 1982). Finding its peptides in PSI-containing bands and in PSII-containing bands of the stroma series is the expected distribution, and on this gel the peptides are spread across the series rather than nested inside PSI (Table S1). The *stn7* high-mass bands 1 and 4 carry a larger fraction of the band-10 maximum (0.55 and 0.43) than the *pph1* bands 2 and 3 (0.30 and 0.32); while band 8 is the same in both genotypes. We also observe elevated bands 1 and 4 in stn7. This is not yet a genotype-specific cytochrome particle in this molecular-mass region of the gel from which the band is isolated.

The free-flow separation asks a different question and still does not name a particle. Two proteins in the same well share a charge window (Fig. 4). Clustering of wells 30–60 then asks whether their charge profiles have the same shape (Fig. S3A). Lhcb1 and Lhcb2 show a joint bimodal distribution at the anodic and cathodic fractions of the separation range; the minor antenna is anodic and tracks PSII, and cytochrome *b*_6_*f* remains its own bar throughout. Notably, complexes that run farthest toward the anode in free solution are among those the gel retards most. In both separation techniques, charge drives the complexes’ mobility; however, a shared charge window does not mean a shared molecular weight, and retardation/filtration upon entry into the gel resolves this.

On this charge axis, the PSI bar of *stn7* peaks at well 58, cathodic of the NDH bar at well 46, and the PSI bar of *pph1* peaks at well 46, in the same well as NDH (Fig. S3A). CURT1A follows the PSI profile more closely in *pph1* (*r* = 0.86), where the leaf sits beside PsaG, than in *stn7* (*r* = 0.70), and the leaf remains outside the PSI bar in both genotypes (Fig. S3B and Fig. S3C). AT5G02160 remains outside the ATPase bar and occupies the anodic window shared by PSII and ATP synthase. A shared well on this axis is a charge window. The particle identified here is the complex of PSI–LHCI with cytochrome *b*_6_*f* as resolved by analysis the second native gel.

The second native dimension we applied after 1D iZ-FFE restores the in-gel molecular weight analytical band structure. The same wells, re-run on LN-PAGE, separate a 720 kDa class in both genotypes from a 753 kDa class that is *pph1*-specific (Figure 5A). The 753 kDa class is more anodic than the 720 kDa class (Figure 5B). Phosphate on the mobile LHCII is the expected source of that anodic shift, and the extra apparent mass is LHCII1,2, placed at band 7 rather than band 8 in the 1D LN-PAGE separation (Fig. 2A). In the MS analysis that we restricted to those fluorescent, and coomassie stainable gel slices of equal molecular weight, both 720 kDa trees are PSI–LHCI–Cyt *b*_6_*f*, with PetA and PetB inside the PSI–LHCI node (*stn7 d* = 0.50; *pph1 d* = 0.34; Table 1). The *pph1* 753 kDa class is a different mixed node (*d* = 0.38). On the tree drawn at a per-slice maximum of 10, PetA and PetC are leaves with Lhcb1.2, Lhcb1.5, and Lhcb2.2, but PetB is not. Across the five slices of that mass class, PetB is present in every slice (5, 6, 6, 6 and 6; sum 29) but based on our cut off definition, it is not a leaf. PetA (sum 44) and PetC (sum 48) are leaves, together with Lhcb1.2, Lhcb1.5 and Lhcb2.2. Table 1 reports the collapsed composition. The tree of the 753 kDa class is named PSI–LHCI–LHCII1,2. PetB is not a leaf of that tree, and the summed counts do not represent a copy number. Band 7 is the larger, more anodic class relative to the PSI–LHCI–cytochrome *b*_6_*f* complex identified at 720 kDa.

We set a floor of 10 counts for accessory proteins to be included in the unsupervised HCA tree. Structural databases do not regard accessory proteins as subunits of a protein complex particle. We included them because a co-migrating trace can indicate a protein involved in regulating metabolism, assembly, chlorophyll binding to a chlorophyll-binding photosystem, kinase-phosphatase pairs, or excitation and electron flow. Sparklines were the supervised fingerprint analogy to the hierarchical similarity approach and were used to notice such a protein when it is usually lost. We excluded sputtered proteins (i.e., those absent from many fractions), and if their maximum was low, we did not interpret a sparse identification as a partner. We apply that filter to every unassigned protein. However, we judge a subunit only if the structural database already assigns it to the complex under test. Peptides differ in detection probability, and hydrophobicity, length, and charge state change that probability; and data-dependent acquisition records a sample of the ions present rather than every ion (Liu et al., 2004; Nielsen et al., 2004). However, while spectral counts follow relative abundance over roughly two orders of magnitude (Liu et al., 2004), spectral counts and integrated peak areas agree on which proteins change, but peak areas provide a more accurate ratio (Old et al., 2005). A longer polypeptide yields more peptides than a short one present at the same molar amount, which is why spectral counts are sometimes divided by protein length (Zybailov et al., 2006). A spectrum dominated by peptides from ATP synthase, or photosystem I, therefore reports how readily those peptides were recorded. It does not report that more of that complex was present than of cytochrome *b*_6_*f*. This paper did not length-normalize the counts or convert them to stoichiometry.

In complexome profiling, migration profiles are clustered and compared to reference-complex clusters, so a known subunit is seen with its complex (Heide et al., 2012; Giese et al., 2015; Wittig and Malacarne, 2021). Researchers have scored recovery of reference complexes when half of the annotated pairs in a complex are retrieved (Havugimana et al., 2012). Later complexome workflows added a false-discovery estimate that plain hierarchical clustering does not carry (Amedei et al., 2024). None of these studies sum the slices of one native-PAGE mass class, cut across one free-flow charge zone, and then apply a 10-count limit. This paper includes that step, but it is limited to the second-dimension windows of one apparent mass (§4.6.2).

A complex separated in the flowing buffer film occupies a zone of finite width (Eichacker and Weber, 2026). Here, we treat the zone as a single charge distribution, highest at its maximum and continuous into the fractions on either side, as a Gaussian would describe. We did not fit a Gaussian, and the slices are repeated samplings of one mass class after the second gel has removed other masses. They are not independent biological replicates. In these separations, ATP synthase is the narrowest zone, with its maximum at well 40, and we take that width as the practical resolution along the charge axis, based on direct observation from this dataset.

A single band from one-dimensional LN-PAGE cannot be summed in this way, because the class is represented by one slice only. Complexes that do not pass the pores at the top of the gradient remain at the gel surface (Behrens et al., 2013; Eichacker et al., 2015; Wittig et al., 2006). We interpret this as anodic high-mass material, much of it vesicular, reaching the surface pores first, along with the complexes. This is the gel-entry phase, when complex concentration and pore-crowding are at their maximum, and it appears to remove partners bound on the outer face of the PSI core particle. The one-dimensional and two-dimensional band composition of the preseparated charge-hull therefore differs. The fact that the molecular mass of the bands appears not to have changed considerably indicates that in-gel separation is not based on molecular mass but on hydrodynamic minimum-diameter screening.

Two limits of scope belong with that identification. Lhca2 peptides are missing from some second-dimension slices, which lowers an LHCI-to-PSI ratio taken from counts and does not mean that an LHCI subunit has left the particle. PsbS remains below the clustering floor and is not part of the particle identified here (Li et al., 2000; Betterle et al., 2009). The separations do not recover the *Chlamydomonas* PSI–LHCI–LHCII–cytochrome *b*_6_*f*–FNR– PGRL1 particle as one second-dimension class (Iwai et al., 2010): FNR is spread across the first-dimension bands, and PGRL1 does not seed a core (Arnold, 2013). Bands 1 and 4 defined after the 1D PAGE separation are not used here to discuss cyclic electron flow.

### 3.3 The mobile polypeptides are Lhcb1.3, Lhcb1.4 and Lhcb2.4, and phosphate changes their address

Among the Lhc gene products, Lhca1–4 are highest with PSI in band 8 at 720 kDa, and Lhcb1–6 are highest in the PSII bands 2 and 3 (Fig. 2A). Bands 1–6 contain Lhca1–5 together with Lhcb1–6. Band 8 lacks the mobile Lhcb1 and Lhcb2 polypeptides that reach their maximum in band 7. On the PSII side, band 2 contains the highest amount of Lhcb1–6 and, together with band 7, reacts with the phosphothreonine antibody (Fig. 1B). Bands 1–6 are high-mass complexes of PSI or PSII with their own antenna, and several of them are baskets of mixed Lhc populations (Fig. 2B). The products that reach their stroma maximum at band 7 (753 kDa) and at band 2, and that stay between 0.13 and 0.26 of that maximum at band 8 (720 kDa), are Lhcb1.3, Lhcb1.4 and Lhcb2.4. The antibody signal for Lhcb1 and Lhcb2 at band 8 is essentially absent (Fig. S1). These three products are the mobile antenna, and they match the LHCII substrates of STN7 and PPH1 (Bellafiore et al., 2005; Shapiguzov et al., 2010). Lhcb3 lacks the N-terminal phosphorylation site and is tightly bound in the PSII complex, with its maximum in band 2 (Standfuss and Kühlbrandt, 2004; Bassi et al., 1988). Lhcb5 and Lhcb6 are minor antennae in the C_2_S_2_M_2_ architecture (Caffarri et al., 2009), so their maxima in bands 2 and 3 are the expected positions.

The three Lhcb4 gene products are outside that mobile pool (Fig. 2A). Lhcb4.1 and Lhcb4.2 peak in bands 2 and 3 and remain above half their maximum in *stn7* bands 1 and 4, both of which also contain PSI. Lhcb4.3 reaches its maximum in band 3, is absent from bands 7 and 8, and does not join the PSII bar on the stroma tree (Fig. S2). CP29 is a monomeric minor antenna (Caffarri et al., 2009). The first-dimension gel does not decide whether the Lhcb4 signal in bands 1 and 4 is CP29 bound to PSI or CP29 inside a PSII complex that shares the basket. Lhca5 is a separate case. Its stroma maximum is *stn7* band 1, with a lower count in band 4 and in both cuts of band 8, and none in *pph1* band 7 (Fig. 2A). Four unique peptides do not meet the tree threshold, so Lhca5 does not enter the clustering. Lhca5 has been found in the NDH–PSI assembly (Peng et al., 2009), which agrees with its highest abundance in band 1.

The antenna of plant PSI is Lhca1–4 (Qin et al., 2015). The maize PSI–LHCI–LHCII structure places one LHCII trimer against PsaO, PsaH, PsaL and PsaA (Pan et al., 2018). Plants that lack Lhcb2 do not perform state transitions, and Lhcb1 and Lhcb2 complement one another (Pietrzykowska et al., 2014). In isolated PSI–LHCII, Lhcb2 is phosphorylated on more than 98% of the protein and phosphorylated Lhcb1 is largely excluded from that supercomplex (Longoni et al., 2015). One phosphorylated Lhcb2 is sufficient for formation of PSI–LHCII (Crepin and Caffarri, 2015). Phosphate is the mark that places this trimer at the dock. It is not the path along which excitation moves. A separate LHCII pool serves PSI after long-term acclimation to growth light, and that pool is not the short-term mobile trimer (Wientjes et al., 2013). The 753 kDa class of *pph1* is the biochemical counterpart of the phosphorylated dock. Yadav et al. (2017) imaged a labile contact of PSI–LHCI with dimeric cytochrome *b*_6_*f* and with LHCII in a *pph1* free-flow fraction. The 720 kDa class extends the cytochrome contact to *stn7*. Lhcb1 and Lhcb2 are absent from that node.

Excised *stn7* band 4 emits at 680 nm and at the long-wavelength maximum of PSI–LHCI (Fig. S6; Arnold, 2013). Band 4 also contains NDH and PSII (Fig. 2B). The two emissions show that both emitters are in the slice. They do not demonstrate energy transfer from LHCII to PSI, and they do not demonstrate a state transition that proceeds without phosphorylation. Phosphorylation changes the address of the mobile antenna. The phosphothreonine blot places that signal in *pph1* band 7, not in band 8 and not in *stn7* (Fig. 1B). The same antenna is the anodic 753 kDa class (Fig. 5A, Fig. 5B). The spectra were not searched for the phosphorylated peptide, so the phosphate is not assigned to one of Lhcb1.3, Lhcb1.4 and Lhcb2.4. PetB is present in every slice of the 753 kDa class (sum 29) and is not a leaf of the tree. The tree remains PSI–LHCI–LHCII1,2 (Table 1). Only two classes were taken through interval-zone electrophoresis and the second-dimension clustering: phosphorylated LHCII1,2 at *pph1* band 7 (753 kDa), and PSI–LHCI without that mobile antenna at band 8 (720 kDa) in both genotypes. Band 4 remains a first-dimension basket.

### 3.4 Stable and labile interaction faces, and one connector complex

We used Digitonin at 16 mM to separate the tightly and loosely appressed grana and stroma membrane regions of the thylakoid membrane. The detergent treatment releases the unstacked stroma lamellae and leaves PSII enriched in the granal membranes in a structure that can be separated by centrifugation or filtration (Andersson and Anderson, 1980). Bands 1–11 and bands 12–23 are successive extracts of those two regions, rather than successive fragments of one complex. In intact grana, the particles present together are C_2_, C_2_S_2_, C_2_S_2_M_2_, and cytochrome *b*_6_*f*. Different detergent treatments remove lipids from these complexes, and the associations detected after LN-PAGE are the structures that survive micellar embedding (Mohamed and Oung, 2026). We consider a contact within one association established only when the same subunits are recovered together after charge separation and the second native gel. The 33 kDa step from 720 to 753 kDa is smaller than the polypeptide mass of one LHCII trimer, so apparent mass in LN-PAGE does not reflect subunit mass/number. In contrast, the anodic shift of the 753 kDa class in iZ-FFE is directly related to the negative charge contributed by phosphate (Fig. 5B).

The granal master is the PSII–LHCII supercomplex in bands 12–15, and this membrane compartment contains no 753/720 kDa PSI pair (Fig. S2). Its stable face is the PSII reaction center together with Lhcb3, CP26, CP24, Lhcb4.1 and Lhcb4.2. In the stroma lamellae, CP26, CP24, Lhcb4.1 and Lhcb4.2 lie inside the PSII bar at *d* = 0.61, and Lhcb3 joins that neighborhood at *d* = 0.91 (Fig. 2A, Table S1, Fig. S2). Its labile face is the peripheral trimer of Lhcb1.3, Lhcb1.4, and Lhcb2.4. Those three products also occupy bands 2, 3, and 11 in the stromal membrane fraction, and they are the Lhcb set that reaches band 7. Loss of this trimer before the minor antenna is the order described for the M-LHCII of C_2_S_2_M_2_ (Caffarri et al., 2009). The stromal master is PSI–LHCI. Lhca1–4 remain with it in both genotypes, at band 8 and again in the 720 kDa class (Qin et al., 2015). The added face is one LHCII trimer on the side of the core opposite the Lhca crescent (Pan et al., 2018). Only *pph1* keeps that face, as determined in the 753 kDa class. In the *stn7* mutant, only the 720 kDa class forms and does not form band 7, so the 720 kDa class in *stn7* is the PSI–LHCI of that genotype. In *pph1*, the 720 kDa band may be PSI that never bound the LHCII trimer, or a 753 kDa particle that lost the trimer. This analysis did not differentiate a solubilization series that separates those two.

The Cytochrome *b*_6_*f* and the ATP synthase are two master complexes of their own. The Cyt b_6_f dimer is identified in band 10, and ATPase in band 9 of the stroma lamellae (Fig. 1). Cyt b_6_f is identified in LN-PAGE using a Cyt b_6_ antibody (Fig. 1B), but using MS the complex is identified throughout all bands in the stroma and in bands 17-21 in the grana membrane (Fig. S2). The charge distribution of Cyt b6f markedly differs from that of ATPase. Like the Cyt b6f complex, ATP synthase shows a defined distribution in band 9 and a multi-band location using MS identification. Its charge-dependent separation shows a single peak with a maximum in well 40 (Fig. 4). Cytochrome peptides in the stroma series show this complex in both grana and stroma lamellae (Anderson, 1982). The 720 kDa class, in which cytochrome subunits join PSI after both separations of the stroma membrane, includes PetA and PetB inside the PSI–LHCI clade (Table 1). In the pph1-specific 753 kDa class, the tree, drawn at a per-slice maximum of 10, does not place PetB under a cytochrome bar. Across the five slices of that class, PetB is present in every slice (sum 29), along with PetA and PetC. Table 1 lists those counts. They do not rename the tree, which remains PSI– LHCI–LHCII1,2.

One identified Lhca gene, Lhca5, has been reported to act as an NDH connector. Its maximum is in the *stn7* band 1, and it is absent from *pph1* band 7 (Fig. 2A). The *Arabidopsis* NDH–PSI structure places one NDH between two PSI–LHCI complexes, with Lhca5 on one side of NDH and Lhca6 on the other (Su et al., 2022). These two linkers are required for supercomplex accumulation (Peng et al., 2009). Band 1 is the only stroma band that contains the NDH subunits, and PPL2 joins those leaves in the HCA tree. NdhU (AT5G21430) lies inside that NDH bar. It is a subunit of the electron-donor subcomplex, and the bar keeps the name NDH (Table S1; Yamamoto et al., 2011). MS sequencing of band 1 also revealed PSI, PSII, and cytochrome *b*_6_*f* (Fig. 2A and B). The Lhca5 count was below the tree threshold, and Lhca6 was not identified in the band series (Fig. 2A). The second native gel was not cut for an NDH core. On the free-flow axis, ATP synthase and NDH occupy wells 38–42 and 44–48 (Fig. 4; Yadav et al., 2017). We conclude that Lhca5 and the linker PPL2 mark an NDH–PSI contact inside band 1, and that band 1 remains a basket of protein complexes of similar size. The other stroma membrane proteins identified, such as FNR, FtsH, VAR1, AOS, and HCF136, occupy bands 9–11 and lie outside these faces. The leaves were harvested under short-day conditions that hold *pph1* in State 2 and leave *stn7* without LHCII phosphorylation. The data therefore do not extend to other regimes of light, water, or nutrients, and they contain no *lhca5* genotype with which the connector function could be tested.

The names 720 kDa and 753 kDa are the NativeMark positions of bands 8 and 7. An earlier reading of the same gel family placed wild-type PSI–LHCI at 647 kDa and PSI–LHCI–LHCII at 721 kDa (Yadav et al., 2017), and the same two bands at about 686 kDa and 743 kDa (Arnold, 2013). The step of about 33 kDa between the names used here is an apparent mass. Bands crowd toward the top of a native gel because the logarithm of Mr scales with migration distance (Wittig et al., 2006). The PSII series from C_2_ to C_2_S_2_M_2_ shows the same crowding (Caffarri et al., 2009; Järvi et al., 2011). Those reports show sieving positions. They do not show a marker-calibrated polypeptide difference equal to one LHCII trimer. The fluorescent ladder above the 720 kDa band in *stn7* well 49 is the same kind of apparent-mass shift, and it is not a ladder of oligomers of the PSI–LHCI–cytochrome *b*_6_*f* particle (Fig. S5).

## 4. Materials and methods

### 4.1 Plant material

*Arabidopsis thaliana* Col-0, *stn7* (Bellafiore et al., 2005) and *pph1-3* (SALK_025713; Shapiguzov et al., 2010; Pribil et al., 2010) were used. Seeds were surface-sterilized in 75% ethanol (5 min), rinsed in 95% ethanol, dried, and sown on Murashige–Skoog agar (Murashige and Skoog, 1962) supplemented with 1.5% sucrose and 0.05% MES. Plates were kept at 4 °C for 2 d in the dark, then transferred to a Percival CU36L5 chamber at 22 °C, short day (8 h light / 16 h dark), 100 µE m⁻² s⁻¹. After 10 d, seedlings were transferred to soil and grown at 24 °C under the same photoperiod and irradiance (Arnold, 2013). For the native-PAGE phenotype, plants were adapted to State 2 (3 h at 100 µE m⁻² s⁻¹) before harvest; State 1 in the thesis comparison was 15 h of darkness (Arnold, 2013). Thylakoids were prepared from four-week-old rosette leaves, modified from Kieselbach et al. (1998) and Fristedt et al. (2009), as described (Arnold et al., 2014; Arnold, 2013). Leaves were homogenized four times for 4 s in ice-cold 25 mM tricine/NaOH pH 7.8, 330 mM sorbitol, 1 mM EDTA, 10 mM KCl, 0.15% (w/v) BSA, and 4 mM sodium ascorbate, then filtered and collected. Lysis and washes contained 10 mM NaF. Membranes were stored in 25 mM tricine/NaOH pH 7.8, 100 mM sorbitol, 5 mM MgCl_2_, 10 mM KCl, 10% (v/v) glycerol at 1 mg Chl ml⁻¹, −80 °C. Chlorophyll was determined in 80% acetone according to Porra et al. (1989).

### 4.2 Sequential solubilization and LN-PAGE

Membranes corresponding to 6–8 µg Chl were solubilized in two steps (Arnold et al., 2014; Arnold, 2013). Step 1: 16 mM digitonin in 80 mM tricine, 15 mM Bis-Tris, 0.002% (w/v) LDS, 20% (v/v) glycerol, 0.01% (w/v) Ponceau S, 10 min at 4 °C; uncleared material was pelleted (13 200 rpm, 10 min, 4 °C). The supernatant is the stroma-lamella-enriched extract (bands 1–11). Step 2: The pellet was extracted with 7.2 mM digitonin plus 7.2 mM *n*-dodecyl-β-D-maltoside in the same buffer for the same time, yielding the grana-enriched extract (bands 12–23). Complexes were separated on native LDS-PAGE 3.5–12% Bis-Tris gels (Arnold et al., 2014) with 80 mM tricine, 15 mM Bis-Tris, 0.002% (w/v) LDS as cathode buffer and 50 mM Bis-Tris pH 7.0 as anode buffer. Chlorophyll fluorescence was recorded on a Typhoon Trio (excitation 633 nm, emission 655–685 nm) or an Odyssey imager (680/700 nm). Total protein was stained with SYPRO Ruby (excitation 532 nm, 580 BP 30). Apparent Mr is Rf versus NativeMark Unstained Protein Standard (Invitrogen: IgM hexamer 1236 kDa, IgM pentamer 1048 kDa, apoferritin 720 and 480 kDa, β-phycoerythrin 242 kDa, lactate dehydrogenase 146 kDa, BSA 66 kDa, soybean trypsin inhibitor 20 kDa) on the same gel family (band 8 = 720 kDa, band 7 = 753 kDa). Immunoblots used rabbit sera against phosphothreonine (Zymed), PsaA, Cyt *b*_6_, Lhca1–3 and Lhcb1–6 (Agrisera), detected with anti-rabbit IgG–HRP (GE Healthcare) and ECL (Arnold, 2013). Transfer was semi-dry (Towbin buffer) or wet (Tris–glycine–SDS–methanol) onto nitrocellulose.

### 4.3 77 K fluorescence

Emission spectra of chloroplasts, digitonin extracts, and excised LN-PAGE bands were recorded at 77 K in a glass dewar in a Fluorolog-3 spectrofluorometer (HORIBA Jobin Yvon), excitation 440 nm (thesis band survey: 435 nm; Arnold, 2013). Gel slices of bands 7 and 8 were normalized at 731 nm. FFE-fraction contours are Origin maps of the same excitation, not Excel 3D plots.

### 4.4 iZE-FFE

For FFE, thawed thylakoids (100 µl ≡ 55 µg Chl) were solubilized in 0.25 M sucrose, 16 mM digitonin for 10 min at 10 °C and cleared at 25 000 *rcf*, 10 min, 10 °C; the residual pellet was extracted in 250 mM sucrose, 8 mM digitonin, 8 mM β-DDM under the same centrifugation (FFE LE1 protocol). Samples were supplemented with SPADNS. Digitonin supernatants of *pph1* (instrument sample S1) and *stn7* (S7) were separated on a standard FFE system (FFE Service GmbH; 0.2 mm spacer; PP60 membranes; 10 °C) in HIBA/BisTris media with sucrose and 0.1% digitonin, 1600 V, displayed 86 mA, 24 April 2014, operator utsu (Eichacker and Weber, 2026). Electrode buffers: anode 100 mM H_2_SO_4_; cathode 100 mM NaOH, 200 mM glycine. Anodic stabilization (inlet 1): 100 mM HCl, 50 mM formic acid, 50 mM HIBA, pH 3.8–4.1 (BisTris). Separation: inlets 2–4 pH 5.4; inlets 5–6 pH 6.2; inlet 7 pH 7.0 plus 5 mM NaCl, all with 0.1% digitonin; inlet 8 pH 7.0 without digitonin. Cathodic stabilization (inlet 9): 150 mM HIBA, 375 mM imidazole. Counter-flow: 250 mM sucrose, 50 mM BisTris, 20 mM AMPSO. Sample entered at inlet 5. Interval: load 1 min 10 s, then 4.0 min high voltage at 40 mL h⁻¹ media flow; elution with the field off into 96-well polyethylene plates (well 1 = anode). We read pH per well (Figure 3A). Fractions were concentrated on Vivaspin 100 kDa filters (180–200 µl to 25 µl, 10 °C) for LN-PAGE, or stored at −20 °C before MS processing.

### 4.5 Mass spectrometry — three jobs, never concatenated

We identified peptides by database search, not by de novo sequencing. We acquired three datasets, and we did not fuse any of them.

**MS after 1D native PAGE (Figures 2A, 2B, Table S1).** Fluorescent bands 1–11 (stroma) and 12–23 (grana) were excised from the 1D native gels and processed (Arnold, 2013). Searches: Mascot against TAIR10 (carbamidomethyl Cys, oxidation Met). The working matrix is unique peptide counts per AGI. Default columns were cut from *pph1-3*; starred columns 1*, 4* and 8* (lab codes 2C, 5C, 9C) and bands were cut from *stn7* (Arnold, 2013, Appendix 1). Wild type was not sequenced. Cut-off for Figure 2B = 2 unique peptides. HCA cutoff: window max ≥ 10.

**MS after liquid FFE (Figure 4 and Figure S3A).** Aliquots of even wells 30–60 were taken from a free-flow separation of the two mutants that is not the 24 April 2014 spectroscopic run and not the 25 November 2013 separation loaded on the second gel (projects 2013-018 and 2013-019). We digested them (modified FASP; Wiśniewski et al., 2009) and measured them by LC-MS as described (Distler et al., 2014). Continuum LC-MS data were processed in ProteinLynx GlobalSERVER 3.0 against UniProt *A. thaliana* (version 2013/03/22) with 3 ppm precursor / 10 ppm product tolerance, carbamidomethyl Cys (fixed) and oxidation Met (variable). Label-free quantification used ISOQuant. Probe7 = *stn7*, Probe8 = *pph1*. Units are fmol-like, **not** unique-peptide counts. Floor for a core in Figure 4 = 1.0. HCA of wells 30–60 (Figure S3A–C) uses a fmol-like window max ≥ 2. Table S1 uses a unique-peptide window maximum of at least 10. Table 1 and Figure 5C use a spectral-count window maximum of at least 10 for the proteins that enter the tree. Section 4.6.2 states when that floor is applied to a sum instead. The three datasets stay separate because the units differ: unique-peptide counts on the first gel, fmol-like intensities in free solution, and spectral counts on the second gel. The supervised reading and the unsupervised reading are two uses of one matrix. They are not two instruments.

**MS after 2D LN-PAGE (Figures 5B and 5C).** Excised 2D bands from LN-PAGE of FFE wells (BN4536 *pph1*, BN4537 *stn7*, experiment 6965, 25 Nov 2013) were destained, reduced (DTT), alkylated (iodoacetamide) and digested with trypsin overnight at 37 °C. Peptides were separated on a nanoAcquity UPLC (Waters; Trap Symmetry C18 180 µm × 20 mm, analytical BEH C18 75 µm × 250 mm, 300 nl min⁻¹, 5–35% acetonitrile in 0.1% formic acid over 60 min) and measured on an LTQ Orbitrap Velos (Thermo; resolution 30 000; Top 20; ions ≥ 2+). Data were processed in Proteome Discoverer 1.2 and searched with Mascot against TAIR10 (carbamidomethyl Cys, oxidation Met). Units are spectral counts. Floor for Figure 5B = 2.0. Proteins enter the HCA matrix at a window maximum of at least 10 spectral counts. The composition of one apparent-mass class is the sum defined in §4.6.2. 44-HMW is a high-Mr cut of well 44, not a slice between 44 and 45; 49-5 is placed once at well 49.

### 4.6 Two readings of a count matrix: a supervised reference-based versus unsupervised HCA analysis

**Supervised — core-complete occupancy (Figures 2A, 2B, 4 and 5B).** Identified proteins were first assigned to complexes using crystal-structure subunit lists and proteomic catalogues: PSII, PDB 3ARC (Umena et al., 2011); PSI–LHCI, 2WSC (Amunts et al., 2010); Cyt *b*_6_*f*, 4H44; NDH (Suorsa et al., 2009); ATP synthase (Binder et al., 1978; Zybailov et al., 2008). For the 1D LN-PAGE analysis that assignment is the sparkline-master table (Arnold, 2013, Appendix 1; Excel sheets Crystal / Master). A slice or well contributed to a complex only if the catalytic core was present (count ≥ floor and ≥ 0.2 of that protein’s own maximum). For Fig. 2A the maximum of each Lhc gene is its maximum inside stroma bands 1–11. For Fig. 2B the maximum is taken on bands 1–23, so an Lhcb protein can have its maximum in the grana and then fall below 0.2 in a stroma band where Fig. 2A still shows it. The cores are: PSI = PsaA+PsaB; Cyt *b*_6_*f* = PetA+PetB (PetC never seeded Cyt); ATPase = AtpA+AtpB; LHCII1,2 ≥ 2 Lhcb1/2; NDH ≥ 2; LHCI ≥ 2. Literature members were then summed and the row was max-normalized among core-ok slices. PsbS, PGR5 and PGRL1 were not averaged into a core (Li et al., 2000). This is a **supervised** summary of occupancy. It does not test whether two cores in the same slice belong to one particle.

**Unsupervised — protein-level HCA (Table S1, Figure S3A, Table 1, Figure 5C).** The same protein × slice (or well) matrix was clustered without using the literature grouping as a constraint: average-linkage Euclidean distance on max-norm traces (Sokal and Michener, 1958). literature_complex coloured leaves **after** clustering. Inner bars mark majority same-complex clades, seeded from the catalytic pairs (PsaA+PsaB, PetA+PetB, AtpA+AtpB, PsbA–D). A Cyt bar requires PetA and PetB in a Cyt-majority node; PetC never seeds Cyt. An outer bar names a mixed occupancy-shape node when those inner clades sit inside it. Windows: stroma bands 1–11 (Table S1; unique-count window max ≥ 10); 1D FFE even wells 30–60 (Figure S3A–C; fmol-like window max ≥ 2); 2D 720 kDa *stn7* columns 46, 47, 48, 49-5, 50, 51, 52 and 53, and *pph1* columns 48-2, 50-2, 51-2 and 52-2; 2D 753 kDa *pph1* columns 43–47 (Table 1 and Figure 5C; spectral-count window max ≥ 10 for the tree; the summed composition is §4.6.2). Column 49-5 is the single 720 kDa slice of well 49. Column 48-2 is the 720 kDa slice of *pph1* well 48 and is not part of the 753 window. Display-only reclustering of those locked windows is allowed; the input matrices were not rebuilt. Euclidean *d* is a shape similarity of traces, not nanometres and not copy number. Sparkline nearest-neighbour of 1D slices (Arnold, 2013) is the historical parent of Reading 1. Reading 1 and Reading 2 answer different questions. Neither replaces the other. The rule that decides where an HCA bar may be drawn, and what that bar is allowed to mean, is §4.6.1.

#### 4.6.1 The distance, the bar, and what may be concluded

Hierarchical clustering is used here for a single quantity. Each protein is a vector of its abundance across the fractions of one separation window, divided by its own maximum in that window. The distance is Euclidean, and the linkage is average (UPGMA; Sokal and Michener, 1958). A small distance means that two max-normalized profiles have a similar shape. It does not mean the proteins are bound, were close in the membrane, or that one copy of one accompanies one copy of the other. The floor used before clustering decides which proteins enter the matrix. The iZE fractions resemble a window maximum of 2 fmol-like units. For gel-band matrices, we applied a window maximum of 10 counts, with unique peptides on the one-dimensional gel and spectral counts on the second-dimension gel. It is an identification threshold. It is not a height at which the tree is cut. Section 4.6.2 notes that the same number is used separately when summing the composition of one second-dimension mass class.

No single height can be called the cutoff for a biologically meaningful interaction, and the co-fractionation literature does not provide one to import. Complexome profiling clusters migration profiles so subunits of a known complex fall together and unassigned proteins with the same profile appear alongside them (Heide et al., 2012; Giese et al., 2015; Wittig and Malacarne, 2021). That literature commonly uses Pearson distance with average linkage, and the profiles are max normalized as here. We interpret the resulting order against reference complexes. A constant height cut on a dendrogram is known to fail when true clusters sit at different heights (Langfelder et al., 2008). When software uses a numerical cut, it is a parameter of that analysis. ComPrAn cuts a Pearson distance at 0.85 (Páleníková et al., 2021). That value is not a constant of protein interaction, and it is not on the Euclidean scale of the trees in this paper. Hierarchical clustering of complexome profiles has since been described as dependent on choices that carry no false-discovery estimate, which is why later workflows add one (Amedei et al., 2024). We therefore do not adopt a universal *d*, Euclidean or otherwise, as evidence that two complexes interact.

The crystallized and atomically analyzed complexes remain the reference sets. For photosystem II, photosystem I, the ATP synthase, the NDH complex and cytochrome *b*_6_*f*, the subunits required for the complex are known from the structures (Umena et al., 2011; Amunts et al., 2010; Stroebel et al., 2003; Suorsa et al., 2009; Hahn et al., 2018). Reading 1 asks whether these structures can answer the following: once the catalytic pair is present, do the remaining structural subunits share a distribution close enough that one supervised trace is a fair summary? The sparkline answers that question, and nothing in the clustering contradicts the subunit lists. What the sparkline does not supply is a reported distance, a test of proteins that were never on the list, or a way to see whether two reference sets join each other before either joins the rest of the matrix. Those are the questions Reading 2 is for.

In this work, a bar on a tree is the lowest node that contains a declared reference pair, and it is drawn only at that node. The reference pairs are PsaA with PsaB, D1 with D2 together with CP43 and CP47, PetA with PetB, and AtpA with AtpB. PetC does not seed cytochrome *b*_6_*f*. The NDH reference is the set used for that complex in the corresponding bar table, and it is stated with the bar. The distance printed on the bar is the height of this node. A join above the bar is a larger set. Its height is reported with the leaves that enter at that join, and it does not replace the distance printed on the bar. On the stroma tree the drawn bars are LHCII1,2 at *d* = 0.19 (Lhcb1.4 and Lhcb2.4; Lhcb1.3 enters at 0.37), PSII at *d* = 0.61 (Lhcb3, PsbS and ascorbate peroxidase 4 enter at 0.91), PSI–LHCI at *d* = 0.56, cytochrome *b*_6_*f* at *d* = 0.71, ATP synthase at *d* = 0.17 (AtpA and AtpB; the seven subunits are together at 0.70) and NDH at *d* = 0.11 (Fig. S2, Table S1). These heights differ between complexes and between windows because the profiles differ. That variation is reported. It is not adjusted to a common cutoff. Every leaf under the node is covered by the bar. The bar carries the reference complex name only when a majority of those leaves belong to that complex in the structure list. Leaves under the node that are not in the list are co-migrating proteins. They are named as such and are not absorbed into the complex. A second bar is drawn inside the first only when it is the node of a second reference set, as when Lhca3 and Lhca4 join before the photosystem I pair does. An outer bar is drawn when two reference pairs both sit inside one node. This means the two reference profiles are more similar to each other than either is to the proteins outside the node.

From that statement, we may conclude co-distribution along the clustered axis. Co-distribution is required for subunits of one particle, but it can also arise from separate particles that share a charge window or a mass window. The particle conclusion is available only when the same two reference sets still join after the material has been separated on the orthogonal axis, within the peptide floor of that matrix. Nesting on one tree is the observation that prompts that test. It is not the test. Section 4.6.2 states the further report that is allowed for a known subunit inside one apparent-mass class of the second gel: the subunit can be part of the summed composition of that class when it does not enter the tree.

On the *pph1* tree of wells 30–60, the node named LHCII contains two CP29 accessions and Lhcb3 (*d* = 0.26, three leaves). CP24 and CP26 are outside it. A profile that averages CP24, CP26, CP29, and Lhcb3 therefore describes a literature group of the minor antenna, which is a Reading 1 trace, and does not describe an HCA clade. The photosystem I node of that tree (*d* = 0.47) already contains Lhca3 and Lhca4; the separate LHCI node (*d* = 0.16) is the inner reference pair. Separate panels for photosystem I and LHCI can be shown, as in Figure 4, when the purpose is to compare two reference sets. They cannot be shown as two HCA complexes discovered independently when one node sits inside the other. The same discipline applies to every bar in Table 1, Table S1, Fig. S2, Figure S3 and Figure 5C.

#### 4.6.2 Where the count of 10 is applied

The value 10 is an identification cut-off. It is applied in two places, and each place answers a different question. An unassigned protein enters a clustering matrix only when its highest count in the window is at least 10 (LN-PAGE) or at least 2 fmol-like units (iZE). The cut-off was set for proteins whose counts are scattered across bands or free-flow fractions and whose maximum is also low. Those two features were treated as technical noise. A protein that fails the floor is not proposed as a new complex partner. This use is unchanged for the stroma tree, the grana tree, the charge trees of wells 30–60, and for every unassigned protein on the second-dimension trees.

A subunit that a crystal structure already assigns to a reference complex is treated differently and is considered only within one apparent-mass class of the second-dimension gel. The slices of that class are one mass, cut from successive fractions along the charge axis, after the second gel has removed complexes of other masses. Their counts are repeated samplings of one class. Table 1 reports each class as a single composition. A known subunit is included in that composition when it is detected across slices and its total count is at least 10. It does not have to reach 10 in any single slice, and it does not have to be a leaf of the tree. The tree is still built at the per-slice floor. A subunit that never reaches 10 in one slice is absent from the dendrogram, and no bar is drawn from a reference pair that is not in the matrix.

The sum is not used for a one-dimensional band, for a free-flow well, or for slices that do not share one apparent mass. The five slices of *stn7* well 49 are not one mass class: 49-5 is the 720 kDa band and 49-1 through 49-4 lie above it. A protein that passes the summed floor is not absorbed into a complex it does not belong to. ATP synthase subunits detected inside the 753 kDa window remain ATP synthase subunits. The sum is also not a copy number. Detection depends on peptide length, ionization, and data-dependent sampling (Liu et al., 2004; Nielsen et al., 2004; Old et al., 2005; Zybailov et al., 2006). No ratio of cytochrome *b*_6_*f* to photosystem I is calculated. In the *pph1* 753 kDa window (wells 43–47), the spectral counts of cytochrome *b*_6_ (PetB) are 5, 6, 6, 6, and 6 (sum 29, maximum 6, five slices out of five). Cytochrome *f* (PetA) sums to 44 and the Rieske protein PetC to 48. PetB does not enter the tree. PetA and PetC do. The collapsed composition includes all three. PetD contributed no counts in this window. Table 1 reports the same counts for each 720 kDa window on its own bands-combined row (PetA 216, PetB 86, PetC 103 and PetD 15 in *stn7*; PetA 98, PetB 40, PetC 51 and PetD 6 in *pph1*). PetB is already a leaf of those trees. The two 720 kDa windows are not added together.

### 4.7 Display conventions

*stn7* solid filled, *pph1* dashed open. Both cuts of gel band 8 are labeled **8**. 44-HMW omitted from charge series. 49-5 once, at well 49. Helper lines: solid grey 0.5 and 1.0 behind the data, dashed 0.2 in front. FFE *x*: anode left, cathode right. Complex order top-down: ATPase, NDH, PSI, LHCI, Cyt *b*_6_*f*, LHCII1,2, PSII.

## Acknowledgements

We thank Andy Christoforou (Cambridge Centre for Proteomics) for discussions of hierarchical clustering, and Ute Sukop-Köppel (FFE Service GmbH) for operating the free-flow electrophoresis. We thank Stefan Tenzer (Johannes Gutenberg-Universität Mainz) for mass spectrometry of the free-flow fractions. We thank Jean-David Rochaix and Michel P. Goldschmidt-Clermont (University of Geneva) for the *stn7* and *pph1* lines and for discussions during the Geneva cooperation. The interval-zone free-flow method is described separately (Eichacker and Weber, 2026). This work was supported by the Research Council of Norway (project 335017).

## CRediT authorship contribution statement

Lutz A. Eichacker: Conceptualization, Investigation, Formal analysis, Writing – original draft, Supervision, Funding acquisition. Janine Arnold: Investigation. Geoffrey Fucile: Resources. Ann Kristin Vatland: Investigation. Karol Stensen: Investigation. Gerhard Weber: Methodology, Resources.

## Data availability

The identification lists and the raw LC-MS files that support this article are kept under the rule stated in the final data-management plan filed with the Research Council of Norway for project 335017 (31 August 2026). Primary instrument files remain with the laboratory that recorded them. The subset used for this manuscript is held on the corresponding author’s University of Stavanger computer and on UiS institutional storage (OneDrive / Microsoft 365). The University of Stavanger keeps project data delivered to UiS for at least ten years after the project period, which ended on 30 June 2026. No DataverseNO dataset, and no other repository accession, has been issued for these files. A selected dataset can still be archived before 30 June 2028.

## Figures and supplementary figures

Figures prepared for Biochimica et Biophysica Acta — Bioenergetics

## Separate supplement file

Supplementary table and figures are in Figures_Supplement_2.pptx (Table S1 and Figs. S1–S6).

