## Supplemental table S1, and figures S1-6 for "Mass spectrometric identification of a PSI–LHCI–cytochrome *b*_6_*f* complex in both *Arabidopsis stn7* and *pph1*"

The supplement contains the hierarchical clustering that tests the sparkline reading, and the gels and spectra that document the separations. On the stroma gel and on the free-flow axis, the tree confirms the sparkline and tightens it. The identification of the PSI–LHCI–cytochrome $b_6f$ particle is in the main text (Fig. 5C, Table 1).

#### Supplementary table and figure legends

**Table S1.** Hierarchical clustering of stroma native-PAGE proteins (bands 1–11; unique-peptide window maximum at least 10; full tree Fig. S2). A bar is the node drawn in Fig. S2, and *d* is the height of that node. Every leaf under the bar is inside that *d*. A higher join is named with the leaves that enter there. Band numbers follow Fig. 1A. IgG is the antibody result for PsaA, Cyt $b_6$ and phosphothreonine (Fig. 1B) and for Lhca and Lhcb (Fig. S1). Three rows are not assigned to a bar. NdhU (AT5G21430; CRRL) inside the NDH bar ($d$ = 0.11) is a J-domain subunit of the NDH electron-donor subcomplex (Yamamoto et al., 2011; Ifuku et al., 2011). The LHCII1,2 bar is Lhcb1.4 and Lhcb2.4 at $d$ = 0.19; Lhcb1.3 joins at 0.37. The PSII bar is $d$ = 0.61; Lhcb3, PsbS and ascorbate peroxidase 4 join at 0.91. The ATPase bar is AtpA and AtpB at $d$ = 0.17. AtpD and AtpE join them at 0.34, AtpC joins at 0.47, and F0 B' and AtpF join at 0.70.

**Fig. S1.** Immunoblot of Lhca (A1–A3) and Lhcb (B1–B6) on the same native LN-PAGE family as Figure 1B. The 1D LN-PAGE lane of pph1-3 is on the left and the lane separating the stroma lamellae thylakoid membrane protein of stn7 is on the right. Gel strips on the top are showing chlorophyll fluorescence of stroma bands labelled 1–11. Lhca (A1-A3) track the PSI complex with highest signal strength at bands 7 and 8 in pph1 and stn7. The Lhcb1 and Lhcb2 (B1, B2, dashed box) gel-blots identify band 2,3, 7, and 11, and are essentially absent from band 8 in pph1. In stn7, these two proteins are specific in band 2, and 4 and the LHCII trimer band 11. The Lhcb (B1-B6) and especially B4 and B6 label PSII complexes 2, and 3 in pph1, and bands 2, and 4 in stn7, and indicate the presence of a LHCII complex in band 10. In combination with Lhca which defines the band distribution of PSI, Lhcb proteins label the bands with mixed PSII and PSI presence, especially in band 1 and 4 in stn7. The result from this blot is shown in the IgG column of Table S1 and resembles an independent experimental control to the MS identifications shown in Figure 2A.

**Fig. S2.** Protein-level average-linkage Euclidean HCA of stroma and grana proteins after LN-PAGE, on the same max-normalised unique-peptide matrix as Fig. 2 (window maximum at least 10). (A) Stroma bands 1–11 (n = 66), summarised in Table S1. The distance cited for a complex is the height of the bar drawn on this panel. A higher join is named separately and lies outside that bar. (B) Grana bands 12–23 (n = 60). On the grana tree the cytochrome bar (PetA, PetC and PetB, *d* = 0.70) sits inside the ATPase bar (*d* = 0.93). The bar is named ATPase. The shared node is co-migration at 441 kDa, not a particle. The LHCII1,2 bar is Lhcb1.3 and Lhcb2.4 (*d* = 0.29). PSI–LHCI is its own bar and does not join cytochrome $b_6f$ or LHCII1,2, so this compartment does not contain the 720 kDa or the 753 kDa particle.

**Figure S3A.** Charge profiles of the bars on the wells 30–60 trees (Fig. S3B and Fig. S3C). Each strip is the mean of the max-normalised traces under that genotype's bar, then scaled to its own maximum (fmol-like floor 2). The order is ATPase, NDH, PSI, LHCI, cytochrome $b_6f$, LHCII1,2, PSII, CP29–Lhcb3 and FNRL2. The PSI strip is the photosystem I node, which already contains the LHCI pair; the LHCI strip is that inner Lhca3 and Lhca4 pair. Cytochrome $b_6f$ is its own bar (PetA and PetB, *d* = 0.20 in *stn7*; PetA, PetB and PetD, *d* = 0.38 in *pph1*) and is not drawn joined to PSI. CP29–Lhcb3 is two CP29 accessions and Lhcb3; CP24 and CP26 are outside it. LHCII1,2 is the Lhcb1 and Lhcb2 pair. FNRL2 is the single protein outside that pair and is its own trace. Solid filled symbols, *stn7*; dashed open symbols, *pph1*. The anode is to the left.

**Fig. S3B.** Heatmap of the *stn7* tree drawn in Fig. S3A (even wells 30–60, anode at the left). Each row is one protein in leaf order. Colour is the max-normalised label-free intensity. Cytochrome $b_6f$ is its own bar (PetA and PetB, *d* = 0.20). A higher join with PSI (*d* = 0.70, 17 leaves) is not drawn. The NDH bar includes CEST (AT5G44650), which is not an NDH subunit. The same bar also covers HCF136, which is not an NDH subunit, and the NDH subunits NdhU, PnsL5 (F4K2G0; CYP20-2) and PnsL4 (F4JW56; FKBP16-2) (Ifuku et al., 2011; Yamamoto et al., 2011). CURT1A (AT4G01150) lies outside the PSI bar. AT5G02160 and rhodanese lie outside the ATPase bar.

**Fig. S3C.** Heatmap of the *pph1* tree drawn in Fig. S3A (even wells 30–60, anode at the left). Each row is one protein in leaf order. Colour is the max-normalised label-free intensity. Cytochrome $b_6f$ is its own bar (PetA, PetB and PetD, *d* = 0.38). PetC is outside that bar. The NDH bar is six NDH-family leaves plus CEST (AT5G44650). CEST is not an NDH subunit. CURT1A (AT4G01150) lies beside PsaG and outside the PSI bar. AT5G02160 lies beside PsbO-1 and outside the ATPase bar.

**Figure S4.** Coomassie stain of 2D iZE-FFE/LN-PAGE of mutants stn7 and pph1. Upon iZE-FFE separation, fractions 20 to 60 of mutant plants stn7 (top 2D gel) and pph1 (bottom 2D gel), were concentrated and protein complexes separated by LN-PAGE. The native gel was scanned for fluorescence (Fig. 5A) and thereafter stained using colloidal Coomassie (methods). Protein bands indicated by the black arrowheads inside the gel (white boxes in Fig. 5A) were selected for MS sequencing analysis. A horizontal series of protein complexes referring to a PSI band at 720 kDa (stn7) was assigned to PSI-LHCI-Cyt b6f complexes. The 720 kDa band of pph1 was also assigned to the same type of PSI complex, while the 753 kDa complex in pph1 was assigned to the PSI-LHCI-LHCII1,2 complex by peptide MS analysis and HCA (Fig. 2, Fig. 4, Fig. 5, Fig. S2 and Fig. S3). The corresponding protein structure assignments follow the analysis by Yadav et al., 2017. The cartoons placed under the gels are those structure models. The bands correlated with band 7 (753 kDa) and band 8 (720 kDa) are those assigned by 1D LN-PAGE (Fig. 1A and B). FFE fractions represent charge windows from the anode (left side, low number) to the cathode (right side, high number).

**Fig. S5.** *stn7* well 49, slices 49-5 (the 720 kDa band) through 49-1. Average-linkage Euclidean clustering of max-normalised spectral counts (window maximum at least 10). The PSI bar is PsaA and PsaB with PsaD-1, PsaE-2 and Lhca1 ($d$ = 0.28). Lhca3 is a leaf outside that bar. Lhca4 is below the floor, so there is no Lhca3–Lhca4 pair and no LHCI bar. PetA and PetC are leaves. PetB is below the floor, so there is no cytochrome bar. ATP synthase (AtpA, AtpB and AtpC) is a separate bar ($d$ = 0.34) and peaks in slice 49-1. Slice 49-5 is also a column of Fig. 5C. The five slices are not one mass and are not summed.

**Fig. S6.** 77 K fluorescence emission spectra of fluorescent bands from LN-PAGE (Fig. 1A). Thylakoid membranes of stn7 and pph1-3 were solubilized in two steps and separated by native LDS-PAGE (Fig. 1). For 77 K fluorescence emission spectroscopy, bands representing protein complexes were cut from the stroma-lamellae lane of stn7 (see Fig. 1A) (NDH* (Band 1), PSII-LHCII (Bands 2, 3), mixed (Band 4), or from the grana lane of pph1-3 LHCII (Band 22)) and 77 K emission spectra were recorded from 650 to 800 nm using 435 nm for excitation.

### Figures and supplementary figures

Prepared for Biochimica et Biophysica Acta — Bioenergetics

##### This supplement

Table S1   Hierarchical clustering of the stroma native-PAGE proteins.
Fig. S1   Lhca/Lhcb immunoblots.
Fig. S2   Hierarchical clustering of stroma and grana native-PAGE proteins
Fig. S3   Charge-axis clustering: collapsed profiles (A); stn7 heatmap (B); pph1 heatmap (C).
Fig. S4   Colloidal coomassie stain of the second-dimension gels.
Fig. S5   stn7 well 49, slices 49-5 through 49-1.
Fig. S6   77 K of selected 1D LN-PAGE bands.

##### Main figure file

Main figures and Table 1 are in Figures_Submission_1.pptx.

### Table S1. Hierarchical clustering of stroma native-PAGE proteins

Table S1.Hierarchical clustering of stroma native-PAGE proteins (bands 1–11; unique-peptide window maximum at least 10; full tree Fig. S2). A bar is the node drawn in Fig. S2, and d is the height of that node. Every leaf under the bar is inside that d. A higher join is named with the leaves that enter there. Band numbers follow Fig. 1A. IgG is the antibody result for PsaA, Cyt $b_6$ and phosphothreonine (Fig. 1B) and for Lhca and Lhcb (Fig. S1). Three rows are not assigned to a bar. NdhU (AT5G21430; CRRL) inside the NDH bar (d = 0.11) is an NDH subunit (Yamamoto et al., 2011; Ifuku et al., 2011). The LHCII1,2 bar is Lhcb1.4 and Lhcb2.4 at d = 0.19; Lhcb1.3 joins at 0.37. The PSII bar is d = 0.61; Lhcb3, PsbS and ascorbate peroxidase 4 join at 0.91. The ATPase bar is AtpA and AtpB at d = 0.17. AtpD and AtpE join them at 0.34, AtpC joins at 0.47, and F0 B′ and AtpF join at 0.70.

| Clade | Band (Fig. 2) | d | Leaves under the bar | IgG |
|---|---|---|---|---|
| ATPase | 9 (also 2–8, 10, 11) | 0.17 | AtpB, AtpA. Outside this bar: AtpD, AtpE at 0.34; AtpC at 0.47; F0 B′, AtpF at 0.70 | — |
| NDH | 1 | 0.11 | NdhH, Ndf2, NDH48, NdhM, PPL2, AT5G21430 | — |
| PSI–LHCI | 7, 8 (also 1, 2, 4, 6) | 0.56 | PsaD-1, Lhca3, Lhca4, PsaB, PsaC, PsaF, PsaL, Lhca1, PsaA, Lhca2, PsaH-2, PsaN | PsaA (Fig. 1B); Lhca (Fig. S2A) |
| Cyt b6f | 10 (also 1–9, 11) | 0.71 | PetA, PetC, PetB, Rieske-like | Cyt b6 (Fig. 1B) |
| PSII | 2, 3 (also 1, 4, 5, 8) | 0.61 | CP47, CP26, PsbO-2, CP24, PsbO-1, CP43, CP29, CP29b, D2, D1, TLP18.3 | pThr (Fig. 1B), bands 2–3 |
| LHCII1,2 | 7, 11 (also 1, 2, 3) | 0.19 | Lhcb1.4, Lhcb2.4 | Lhcb1/2 (Fig. S2A); not band 8 |
| not assigned | — | — | pTAC16, LOX2, CaS, PGRL1A | — |
| not assigned | — | — | VAR1, FtsH2, FtsH8, PNDOR, FtsH1 | — |
| not assigned | — | — | AT1G56500, AOS, HCF136 | — |

#### Figure S1. Lhca and Lhcb immunoblots

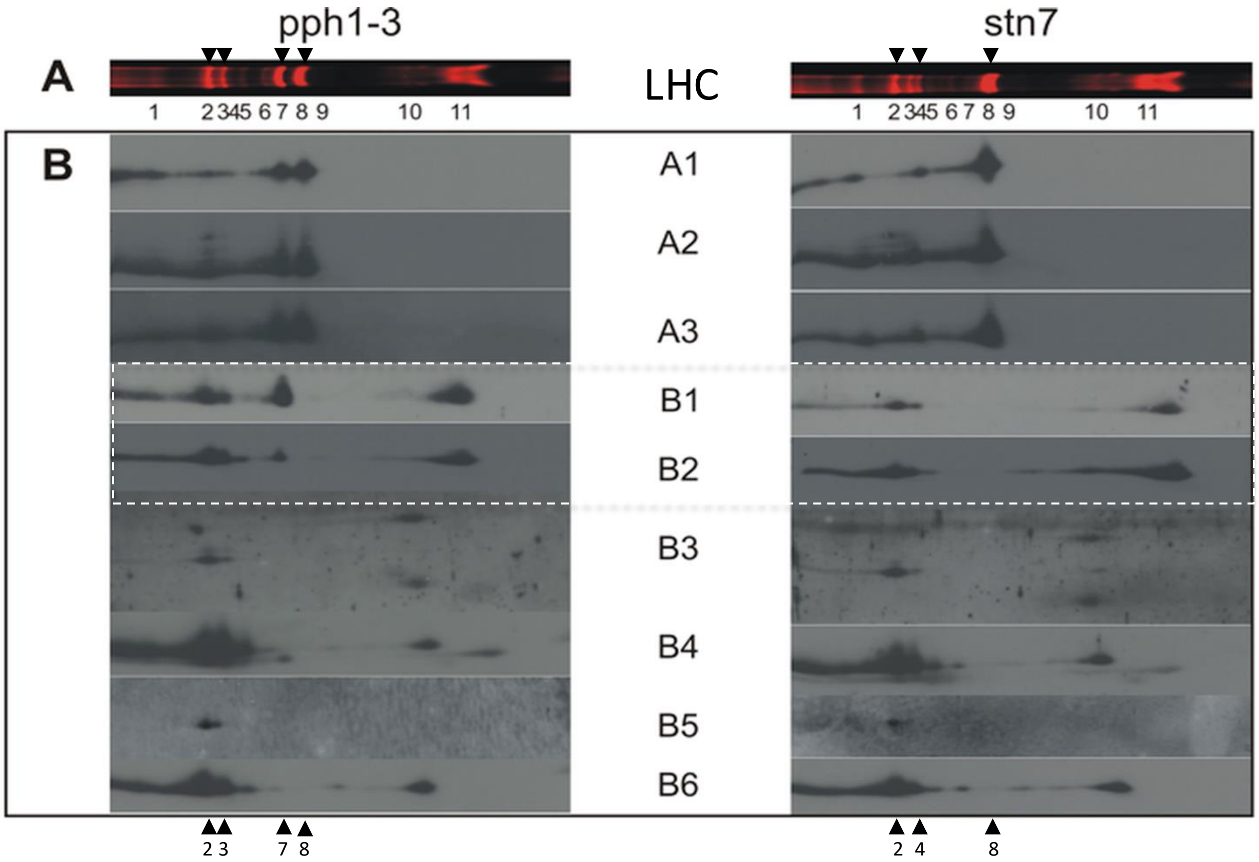


Fig. S1.Immunoblot of Lhca (A1–A3) and Lhcb (B1–B6) on the same native LN-PAGE family as Figure 1B. The 1D LN-PAGE lane of pph1-3 is on the left and the lane separating the stroma lamellae thylakoid membrane protein of stn7 is on the right. Gel strips on the top are showing chlorophyll fluorescence of stroma bands labelled 1–11. Lhca (A1-A3) track the PSI complex with highest signal strength at bands 7 and 8 in pph1 and stn7. The Lhcb1 and Lhcb2 (B1, B2, dashed box) gel-blots identify band 2,3, 7, and 11, and are essentially absent from band 8 in pph1. In stn7, these two proteins are specific in band 2, and 4 and the LHCII trimer band 11. The Lhcb (B1-B6) and especially proteins B4 and B6 proteins label PSII complexes 2, and 3 in pph1, and bands 2, and 4 in stn7, and indicate the presence of a LHCII complex in band 10. In combination with Lhca which defines the band distribution of PSI, Lhcb proteins label the bands with mixed PSII and PSI presence, especially in band 1 and 4 in stn7. The result from this blot is shown in the IgG column of Table S1 and resembles an independent experimental control to the MS identifications shown in Figure 2A.

### Figure S2. Hierarchical clustering of stroma and grana native-PAGE proteins

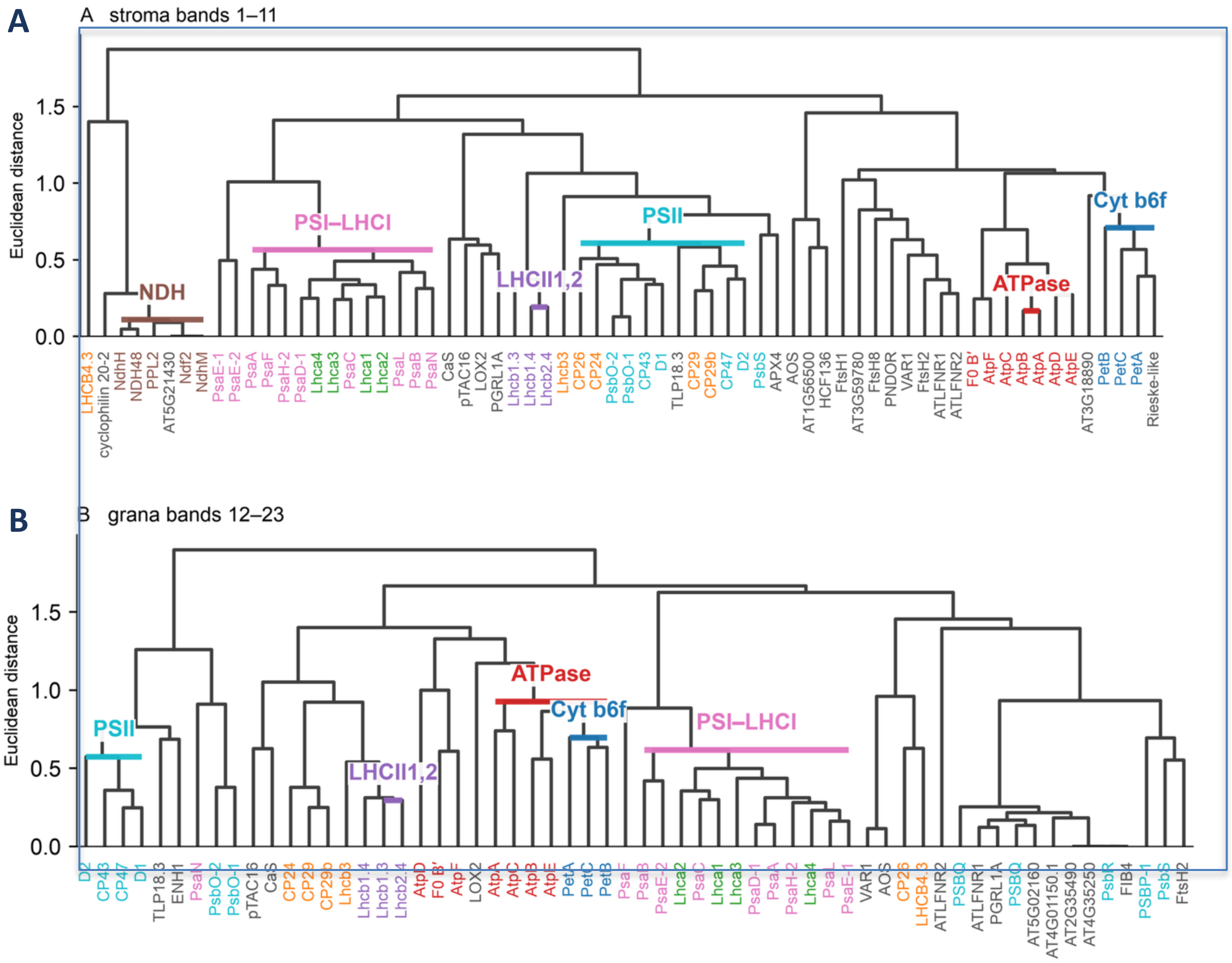


Fig. S2. Protein-level average-linkage Euclidean HCA of stroma and grana proteins after LN-PAGE, on the same max-normalised unique-peptide matrix as Fig. 2 (window maximum at least 10). (A) Stroma bands 1–11 (n = 66), summarised in Table S1. The distance cited for a complex is the height of the bar drawn on this panel. A higher join is named separately and lies outside that bar. (B) Grana bands 12–23 (n = 60). On the grana tree the cytochrome bar (PetA, PetC and PetB, d = 0.70) sits inside the ATPase bar (d = 0.93). The bar is named ATPase. The shared node is co-migration at 441 kDa, not a particle. The LHCII1,2 bar is Lhcb1.3 and Lhcb2.4 (d = 0.29). PSI–LHCI is its own bar and does not join cytochrome $b_6f$ or LHCII1,2, so this compartment does not contain the 720 kDa or the 753 kDa particle.

### Figure S3A. Charge-axis clustering: collapsed profiles (A); stn7 heatmap (B); pph1 heatmap (C).

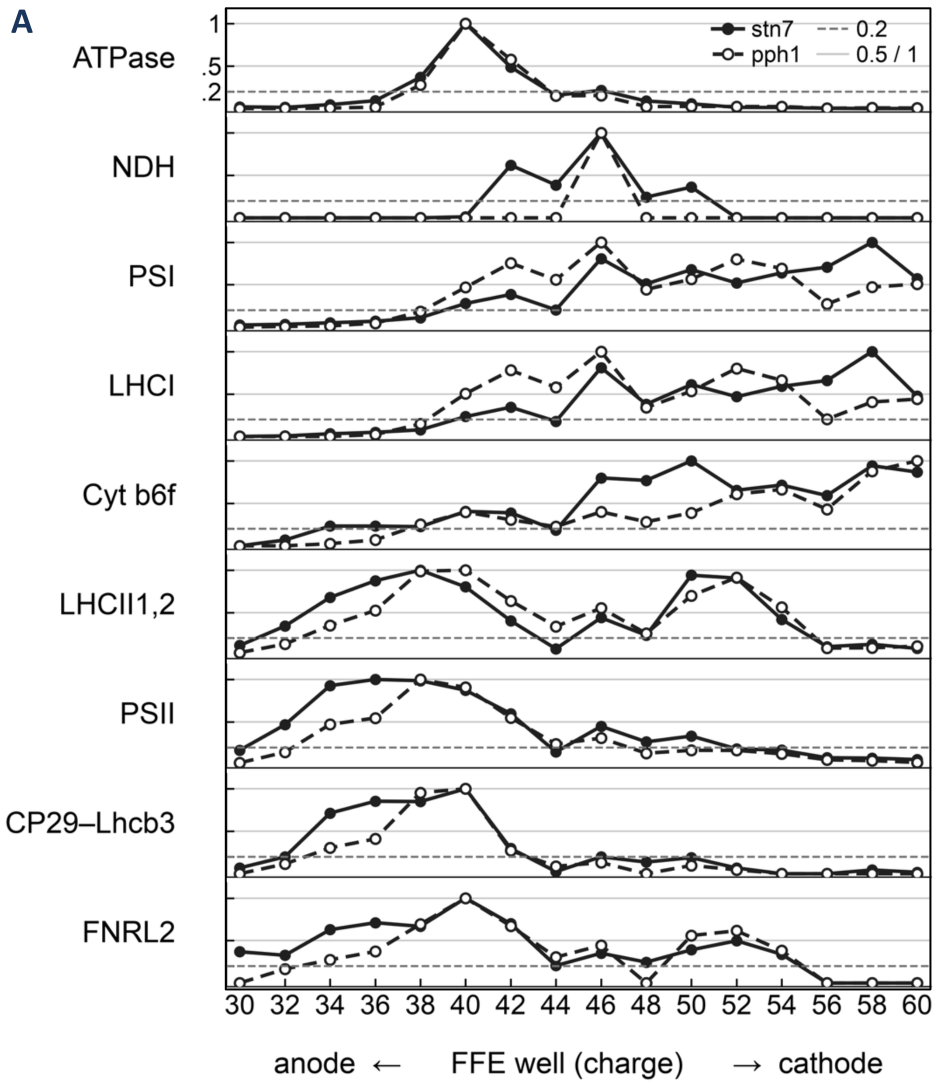


Figure S3A.Charge profiles of the bars on the wells 30–60 trees (Fig. S3B and Fig. S3C). Each strip is the mean of the max-normalised traces under that genotype's bar, then scaled to its own maximum (fmol-like floor 2). The order is ATPase, NDH, PSI, LHCI, cytochrome $b_6f$, LHCII1,2, PSII, CP29–Lhcb3 and FNRL2. The PSI strip is the photosystem I node, which already contains the LHCI pair; the LHCI strip is that inner Lhca3 and Lhca4 pair. Cytochrome $b_6f$ is its own bar (PetA and PetB, d = 0.20 in stn7; PetA, PetB and PetD, d = 0.38 in pph1) and is not drawn joined to PSI. CP29–Lhcb3 is two CP29 accessions and Lhcb3; CP24 and CP26 are outside it. LHCII1,2 is the Lhcb1 and Lhcb2 pair. FNRL2 is the single protein outside that pair and is its own trace. Solid filled symbols, stn7; dashed open symbols, pph1. The anode is to the left.

### Figure S3B. Charge-axis clustering: Heatmap of the stn7 1D FFE tree (wells 30–60)

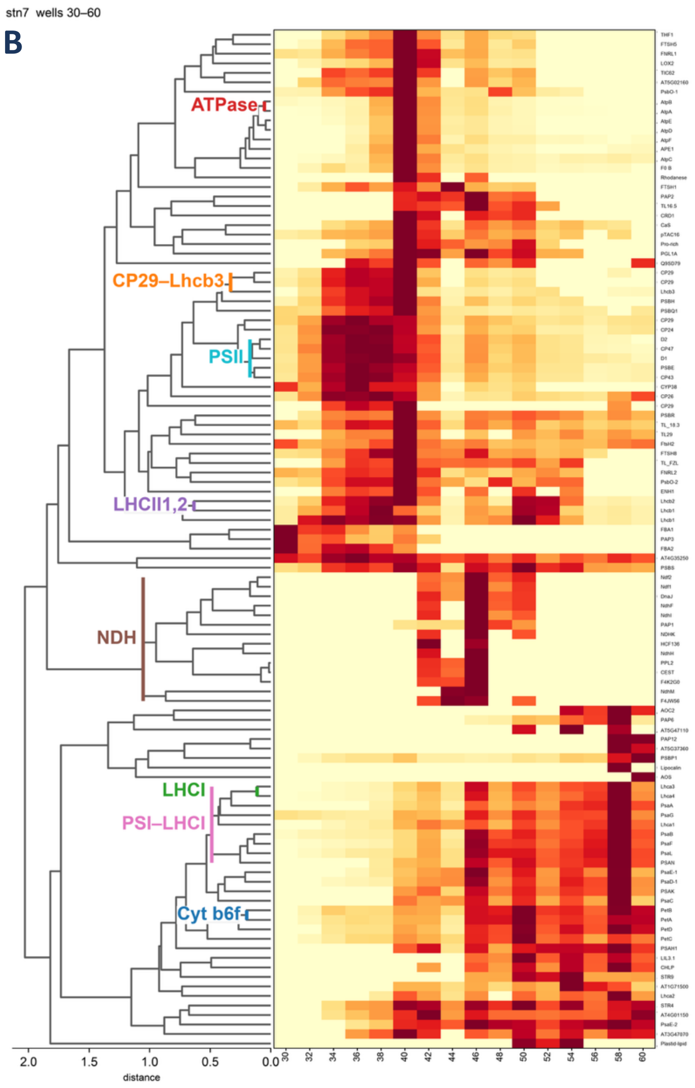


Fig. S3B.Heatmap of the stn7 tree drawn in Fig. S3A (even wells 30–60, anode at the left). Each row is one protein in leaf order. Colour is the max-normalised label-free intensity. Cytochrome $b_6f$ is its own bar (PetA and PetB, d = 0.20). A higher join with PSI (d = 0.70, 17 leaves) is not drawn. The NDH bar includes CEST (AT5G44650), which is not an NDH subunit. The same bar also covers NdhU, PnsL5 (F4K2G0) and PnsL4 (F4JW56), NDH subunits, and HCF136, which is not an NDH subunit. CURT1A (AT4G01150) lies outside the PSI bar. AT5G02160 and rhodanese lie outside the ATPase bar.

### Figure S3C. Heatmap of the pph1 1D FFE tree (wells 30–60)

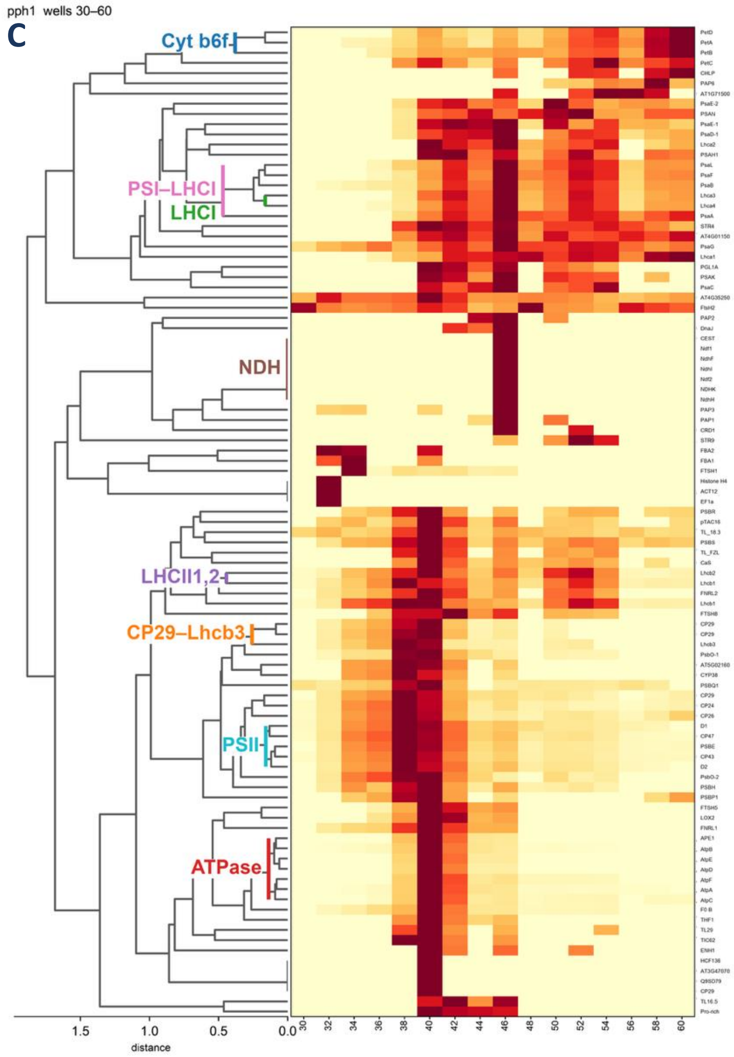


Fig. S3C.Heatmap of the pph1 tree drawn in Fig. S3A (even wells 30–60, anode at the left). Each row is one protein in leaf order. Colour is the max-normalised label-free intensity. Cytochrome $b_6f$ is its own bar (PetA, PetB and PetD, d = 0.38). PetC is outside that bar. The NDH bar is six NDH-family leaves plus CEST (AT5G44650). CEST is not an NDH subunit. CURT1A (AT4G01150) lies beside PsaG and outside the PSI bar. AT5G02160 lies beside PsbO-1 and outside the ATPase bar.

### Fig. S4 Colloidal coomassie stain of the second-dimension gels.

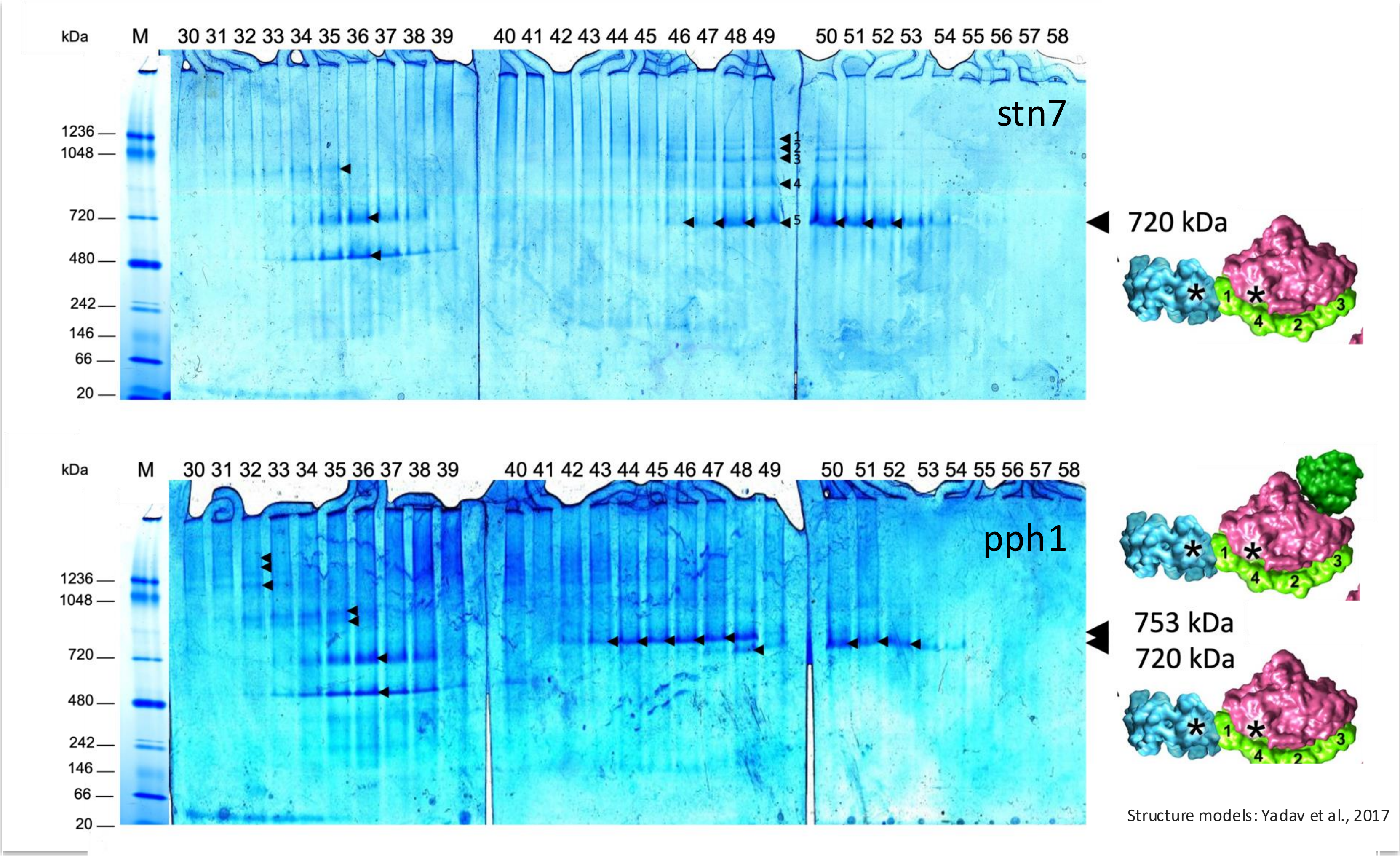


Figure S4. Coomassie stain of 2D iZE-FFE/LN-PAGE of mutants stn7 and pph1. Upon iZE-FFE separation, fractions 20 to 6[illegible] 2D gel) and pph1 (bottom 2D gel), were concentrated and protein complexes separated by LN-PAGE. The native gel was scanned for fluorescence (Fig. 5A) and thereafter stained using colloidal Coomassie (methods). Protein bands indicated by the black arrowheads inside the gel (white boxes in Fig. 5A) were selected for MS sequencing analysis. A horizontal series of protein complexes referring to a PSI band at 720 kDa (stn7) was assigned to PSI-LHCI-Cyt b6f complexes. The 720 kDa band of pph1 was also assigned to the same type of PSI complex, while the 753 kDa complex in pph1 was assigned to the PSI-LHCI-LHCII1,2 complex by peptide MS analysis and HCA (Fig. 2, Fig. 4, Fig. 5, Fig. S2 and Fig. S3). The corresponding protein structure assignments follow the analysis by Yadav et al., 2017. The cartoons placed under the gels are those structure models. The bands correlated with band 7 (753 kDa) and band 8 (720 kDa) are those assigned by 1D LN-PAGE (Fig. 1A and B). FFE fractions represent charge windows from the anode (left side, low number) to the cathode (right side, high number).

#### Figure S5. stn7 well 49 ladder (slices 49-5 to 49-1)

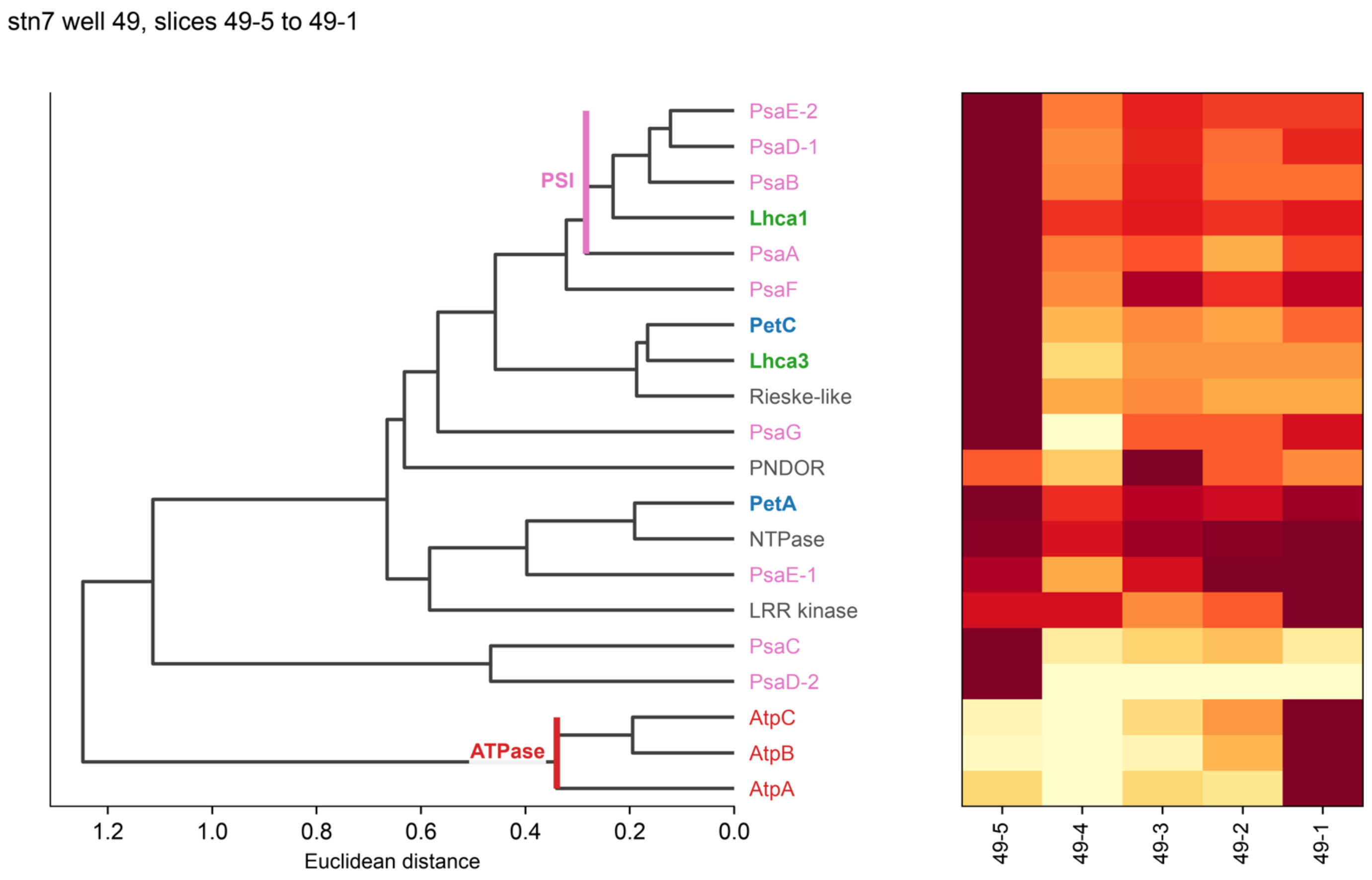


Fig. S5.stn7 well 49, slices 49-5 (the 720 kDa band) through 49-1. Average-linkage Euclidean clustering of max-normalised spectral counts (window maximum at least 10). The PSI bar is PsaA and PsaB with PsaD-1, PsaE-2 and Lhca1 (d = 0.28). Lhca3 is a leaf outside that bar. Lhca4 is below the floor, so there is no Lhca3–Lhca4 pair and no LHCI bar. PetA and PetC are leaves. PetB is below the floor, so there is no cytochrome bar. ATP synthase (AtpA, AtpB and AtpC) is a separate bar (d = 0.34) and peaks in slice 49-1. Slice 49-5 is also a column of Fig. 5C. The five slices are not one mass and are not summed.

# Fig. S6   77 K of selected 1D LN-PAGE bands.

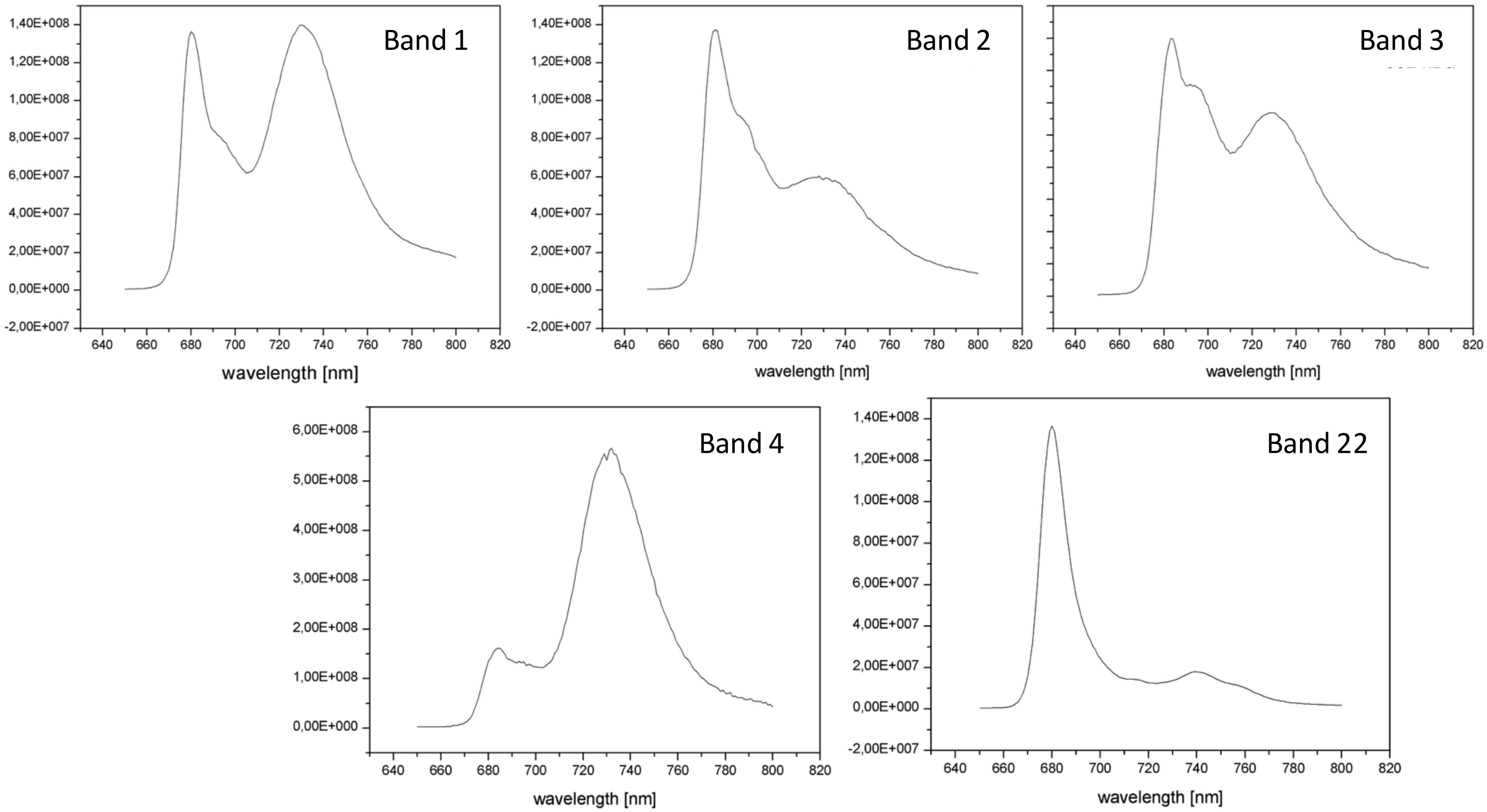


Fig. S6.77 K fluorescence emission spectra of fluorescent bands from LN-PAGE (Fig. 1A). Thylakoid membranes of stn7 and pph1-3 were solubilized in two steps and separated by native LDS-PAGE (Fig. 1). For 77 K fluorescence emission spectroscopy, bands representing protein complexes were cut from the stroma-lamellae lane of stn7 (see Fig. 1A) (NDH* (Band 1), PSII-LHCII (Bands 2, 3), mixed (Band 4), or from the grana lane of pph1-3 LHCII (Band 22)) and 77 K emission spectra were recorded from 650 to 800 nm using 435 nm for excitation.
